# CLEC-2 promotes inflammatory peritoneal macrophage emigration to draining lymph nodes during endotoxemia

**DOI:** 10.1101/2020.12.21.423770

**Authors:** Joshua H. Bourne, Nonantzin Beristain-Covarrubias, Malou Zuidscheroude, Joana Campos, Ying Di, Evelyn Garlick, Martina Colicchia, Lauren V. Terry, Steven G. Thomas, Alexander Brill, Jagadeesh Bayry, Steve P. Watson, Julie Rayes

**Author notes:** **Corresponding authors** Joshua H. Bourne, Institute of Cardiovascular Sciences, College of Medicine and Dentistry, University of Birmingham, Edgbaston, Birmingham B15 2TT, UK, Dr. Julie Rayes, Institute of Cardiovascular Sciences, College of Medicine and Dentistry, University of Birmingham, Edgbaston, Birmingham B15 2TT, UK.

## Abstract

Macrophage recruitment during sterile inflammation and infection is essential to clear pathogens, apoptotic cells and debris. However, persistent macrophage accumulation leads to chronic inflammation. Platelets are emerging as key modulators of the inflammatory response. Here, we identify that platelet C-type-lectin-like receptor-2 (CLEC-2) is a crucial immunomodulatory receptor through the interaction with podoplanin, upregulated on inflammatory macrophages.

Mechanistically, platelet CLEC-2 upregulates the expression of podoplanin and its co-ligands CD44 and ERM proteins, leading to actin rearrangement and promotion of cell migration; this is mimicked by recombinant CLEC-2-Fc (rCLEC-2-Fc). Treatment of LPS-challenged mice with rCLEC-2-Fc induces a rapid emigration of peritoneal macrophages to mesenteric lymph nodes, through a gradient generated by the podoplanin ligand, CCL21, to prime T cells. We propose that crosslinking podoplanin using rCLEC-2-Fc is a novel, cell-specific strategy to accelerate macrophage removal from the site of inflammation, and hence promote the resolution of the inflammatory response.

**Visual Abstract:** 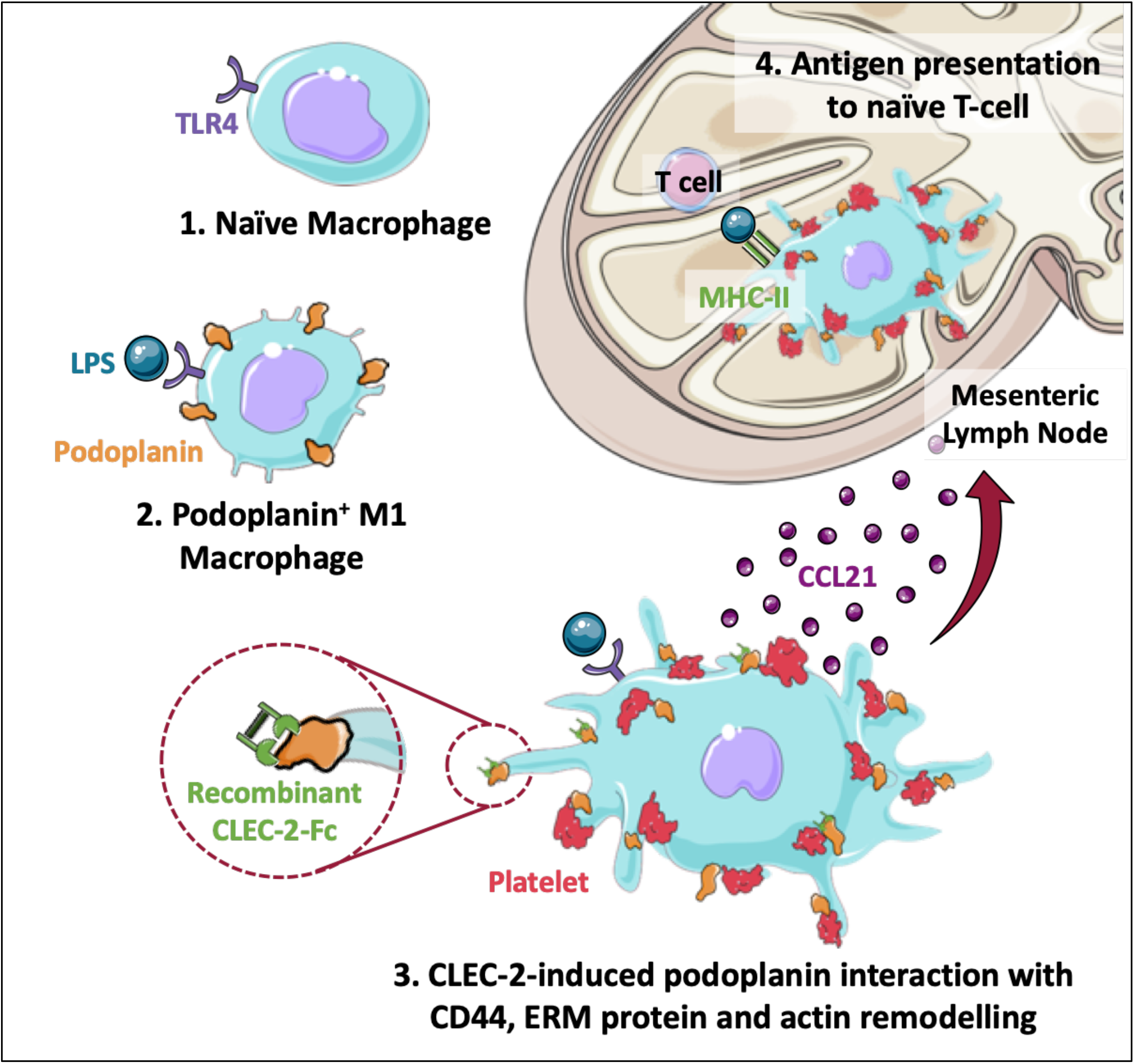

**Summary:** Persistent macrophage accumulation in inflamed tissue leads to chronic inflammation and organ damage. Bourne et al. identify recombinant CLEC-2-Fc crosslinking podoplanin on inflammatory macrophages, as a cell-specific strategy to accelerate their emigration to draining lymph nodes, and reduce local inflammation.

## Introduction

During infection, the development of an adequate inflammatory response is critical to contain pathogen growth and spreading. Innate immune cells are recruited to the site of infection to clear pathogens through multiple mechanisms including the secretion of cytokines and anti-microbial molecules, phagocytosis and through release of extracellular traps (Greenberg and Grinstein, 2002, Kaplan and Radic, 2012). Following pathogen clearance, the resolution of the inflammation is critical to restore tissue homeostasis.

During acute inflammation, from infectious or sterile origins, neutrophils are the first responders (Meng et al., 2012), followed by monocyte mobilisation and subsequent infiltration into the inflammatory site (Italiani and Boraschi, 2014). Monocytes differentiate into inflammatory macrophages with high phagocytic capacity, which secrete anti-microbial molecules, inflammatory cytokines and chemokines. Removal, reprogramming and/or apoptosis of inflammatory macrophages contribute to the resolution of inflammation. A decrease in macrophage death or inability to emigrate from the inflamed site can trigger chronic inflammation (Huang et al., 2009), as seen in many diseases such as atherosclerosis, metabolic disorders or pathogen-induced infections such as SARS-CoV-2. Strategies to accelerate macrophage removal from the site of inflammation constitute a key strategy to promote the resolution of the inflammatory response and reduce organ damage.

Alongside their role in thrombosis and haemostasis, platelets are emerging as vital regulators during the inflammatory response (Rayes et al., 2020). Using different models of acute peritonitis, platelet-macrophage aggregates have been observed in the inflamed peritoneum, but their functional relevance is not well known. During lipopolysaccharide (LPS)-induced endotoxemia, platelets dampen the inflammatory macrophage phenotype, through multiple mechanisms. These include the release of immunomodulatory molecules such as prostaglandin E2 (Xiang et al., 2013) and through the interaction of platelet C-type lectin-like receptor (CLEC-2) with podoplanin upregulated on inflammatory macrophages (Rayes et al., 2017) and monocytes (Xie et al., 2020).

CLEC-2 is a hemi-immunoreceptor tyrosine-based activation motif (hemi-ITAM) receptor constituently expressed on platelets and a sub-set of dendritic cells. Mice with global CLEC-2 deficiency exhibit a lack of lymphatic and vascular endothelial cell separation, which results in blood-filled lymphatics and petechia *in utero* (Bertozzi et al., 2010, Haining et al., 2020), showing a key role for CLEC-2 in the maintenance of vascular integrity during development. CLEC-2 induces platelet activation through its interaction with its endogenous ligands podoplanin (Suzuki-Inoue et al., 2007) or heme (Bourne et al., 2020). Podoplanin is a small, transmembrane O-glycosylated mucin-type protein constituently expressed on type I lung epithelial cells, fibroblastic reticular cells, lymphatic endothelial cells and podocytes (Retzbach et al., 2018, Quintanilla et al., 2019). During cancer and inflammation, podoplanin has been shown to be upregulated on fibroblasts (Farr et al., 1992), Th17 cells (Peters et al., 2011) and macrophages (Kerrigan et al., 2012). Podoplanin promotes epithelial-mesenchymal transition through direct interaction with ezrin, radixin and moesin (ERM) proteins in cancer-associated fibroblasts (Martín-Villar et al., 2006); ERM proteins link membrane proteins to the actin cytoskeleton to induce signalling pathways that regulate cellular motility (Bretscher et al., 2002). However, a role for podoplanin in macrophage migration is yet to be shown.

Beside the role of CLEC-2-podoplanin in thrombosis (Hitchcock et al., 2015, Payne et al., 2017) CLEC-2 regulates the cytokine storm, and decreases the number of peritoneal macrophages (Rayes et al., 2017) and blood monocytes (Xie et al., 2020) during inflammation. Complexes of platelet and podoplanin-positive-macrophage are observed during atherosclerosis (Inoue et al., 2015), rheumatoid arthritis (Takakubo et al., 2017), and breast cancer (Hatzioannou et al., 2016); but how this influences macrophage phenotype and migration is not known.

In this study, we investigate the role of the CLEC-2-podoplanin axis in macrophage functions including phagocytosis, secretion of cytokines and chemokines and their migration. We also assess the functional relevance of this interaction *in vivo* during ongoing peritonitis. Our study demonstrates a key immunoregulatory function for the CLEC-2-podoplanin axis in macrophages, and identifies this interaction as a novel pathway to regulate macrophage trafficking during inflammation.

## Results

### Platelet CLEC-2 upregulates podoplanin and CD44 expression on inflammatory BMDMs and promotes spreading

We have previously shown that platelet CLEC-2 binding to podoplanin, upregulated on the LPS-treated mouse macrophage cell line, RAW264.7, induces platelet aggregation (Kerrigan et al., 2012). We assessed the effect of this interaction on primary mouse macrophages using bone marrow-derived macrophages (BMDMs). LPS induces podoplanin expression on BMDMs isolated from WT (**Fig. 1 A**) but not PDPN^fl/fl^Vav-iCre^+^ mice **(Fig. S1 A)**. Podoplanin upregulation was inhibited by TAK-242 treatment, a TLR-4 specific small molecule inhibitor **(Fig. S1 B)**. Addition of WT platelets for 1h on LPS-stimulated macrophages, but not using CLEC-2-deficitent platelets, further increased podoplanin expression (**Fig. 1 B**) and was associated with an increase in the podoplanin associated membrane ligand CD44, as assessed by flow cytometry **(Fig. 1 C)**. The effect of CLEC-2 is specific to podoplanin and its partner CD44, as for example there was no alteration in CD80 expression **(Fig. S1 C)**. LPS induced a transcriptional upregulation of podoplanin, as demonstrated through podoplanin mRNA upregulation in RAW264.7 cells **(Fig. S2 A)**. However, addition of WT platelets to LPS-activated RAW264.7 cells did not increase the protein level of podoplanin in the whole cell lysate after 1h or 24h **(Fig. S2 B)**. These results suggest that CLEC-2 binding to podoplanin rapidly increases podoplanin and CD44 expression, possibly through the translocation of podoplanin from intracellular stores upon CLEC-2 binding.

**Figure 1:**
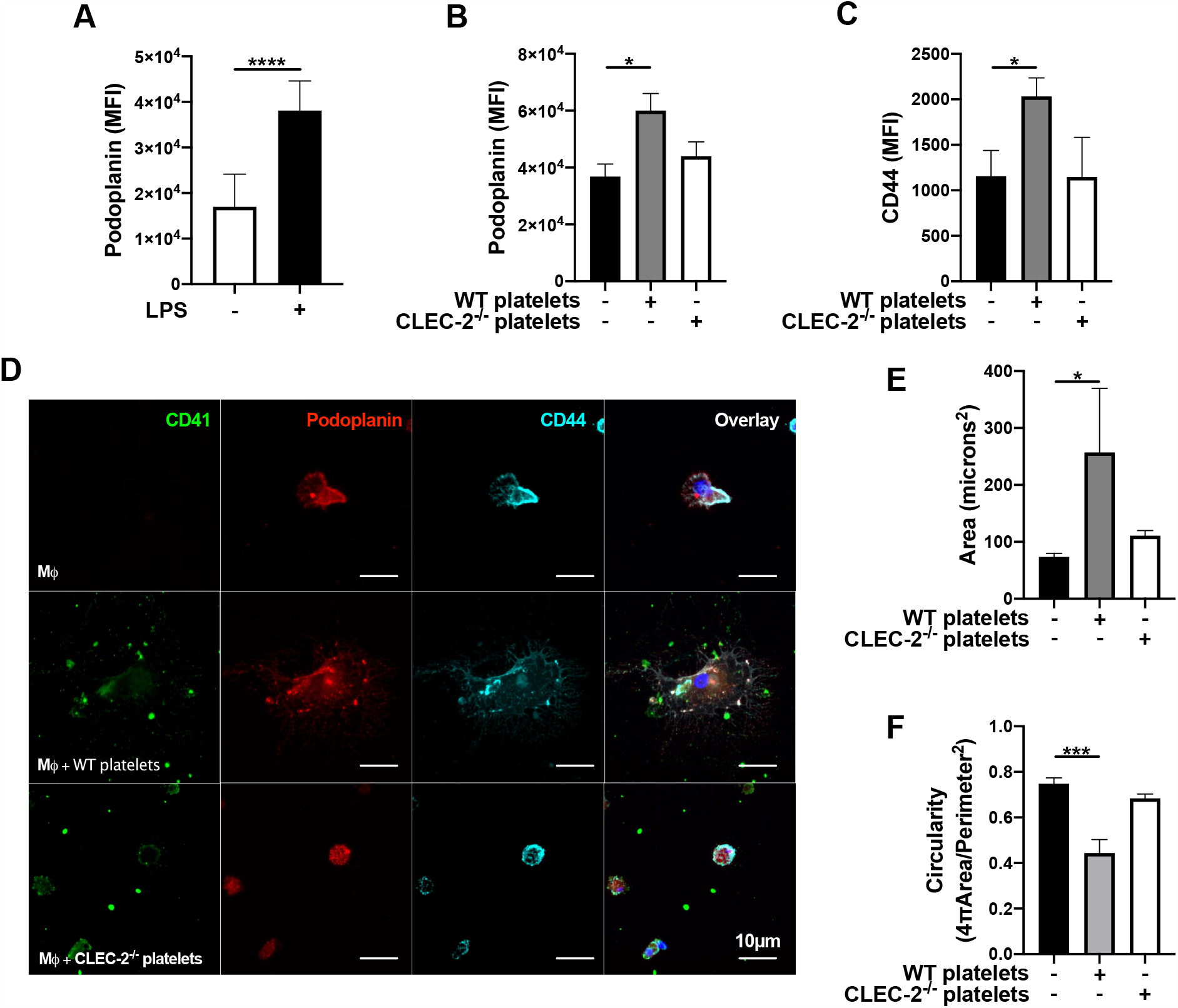
Platelet CLEC-2 upregulates podoplanin and CD44 expression on LPS-stimulated bone marrow-derived macrophages. **(A)** Bone marrow-derived macrophages (BMDMs) were incubated in the presence or absence of LPS (1µg/ml) for 24h. **(B, C)** LPS-stimulated BMDMs (Mϕ) were co-cultured in absence or presence of WT or CLEC-2-deficient platelets (100 platelets: 1 Mϕ ratio) for 1h. **(A-C)** Surface expression of podoplanin (n=4) and CD44 (n=3) median fluorescence intensity (MFI) were detected by flow cytometry using conjugated anti-podoplanin or anti-CD44 antibodies, respectively. **(D)** Mϕ were cultured on glass, and WT or CLEC-2-deficient platelets were added for 1h. Cells were fixed, immunostained for platelets (CD41, green), podoplanin (red) and CD44 (cyan), and detected using confocal microscopy. Images are representative of 4 independent experiments. **(E)** Cell area and **(F)** circularity were analysed using ImageJ. The statistical significance between 2 groups was analyzed using a student’s paired t-test and the statistical difference between multiple groups using one-way ANOVA with Tukey’s multiple comparisons test. \**p*<0.05 \*\*\**p*<0.001 \*\*\*\**p*<0.0001.

We next investigated the translocation of podoplanin from intracellular compartments in inflammatory macrophages in BMDMs and RAW264.7 cells. Macrophages were stimulated with LPS for 24h, and podoplanin expression and its distribution was assessed using immunofluorescence. Following LPS-treatment, podoplanin expression was observed at the pseudopods of both BMDMs and RAW264.7 cells and in granules within the cytoplasm **(Fig. 1 D; S2 C)**. Addition of WT platelets, but not CLEC-2-deficient platelets, increased the spreading of both BMDMs and RAW264.7 cells, as measured by increased cell area and loss of circularity **(Fig. 1 E and F; S2 D and E**), associated with increased pseudopod formation compared to LPS-treated cells.

In order to elucidate the effect of CLEC-2-podoplanin interaction on actin remodelling, LifeAct-GFP-derived BMDMs were assessed using diSPIM LightSheet microscopy. Addition of WT platelets to LPS-activated BMDMs increased pseudopod formation and actin remodelling **(Video 1)** compared to LPS-treated BMDMs **(Video 2)**. In contrast, actin remodelling, spreading and pseudopod formation decreased after the phagocytosis of platelets, showing that the increase in remodelling was independent of uptake. In order to understand whether actin remodelling is associated with an alteration in podoplanin signalling, we assessed the phosphorylation pattern of serines by western blot, as serines 167 and 171 were previously described to be phosphorylated by CDK5 and PKA, and to be associated with a decreased cell mobility (Krishnan et al., 2015). Immunoprecipitation of podoplanin from different co-cultures revealed phosphorylation was inhibited in the presence of WT but not CLEC-2-deficient platelets **(Fig. S2 F)**.

These results show that platelet CLEC-2 binding to podoplanin on inflammatory BMDMs and RAW264.7 cells alters podoplanin downstream signalling, associated with actin remodelling, upregulation of podoplanin and CD44, and their localisation on filopodia.

### Platelet CLEC-2 delays inflammatory BMDM phagocytic capacity and reduces TNF-α secretion *in vitro*

We next investigated whether the modification in cytoskeleton rearrangement alters macrophage phagocytic capacity and their inflammatory phenotype. LPS-stimulated BMDMs were co-cultured in the presence or absence of WT or CLEC-2-deficient platelets and the uptake of pH-sensitive fluorescent *E. Coli*-bound bioparticles was assessed by Incucyte systems for live-cell microscopy **(Fig. 2 A-C)**. Addition of WT platelets significantly delays BMDM uptake of the bioparticles compared to BMDM control. This delay was not observed in the presence of CLEC-2-deficient platelets. Podoplanin deficiency from BMDMs did not affect their phagocytic activity (**Fig. S 3 A and B**) suggesting a distinct role for podoplanin crosslinking by CLEC-2. LPS induced an increase in TNF-α secretion from BMDMs, which was significantly decreased in the presence of WT but not CLEC-2-deficient platelets (**Fig. 2 D**). However, whilst LPS induced an increase in anti-inflammatory IL-10 secretion, WT platelets did not modify IL-10 secretion **(Fig. 2 E)**.

**Figure 2:**
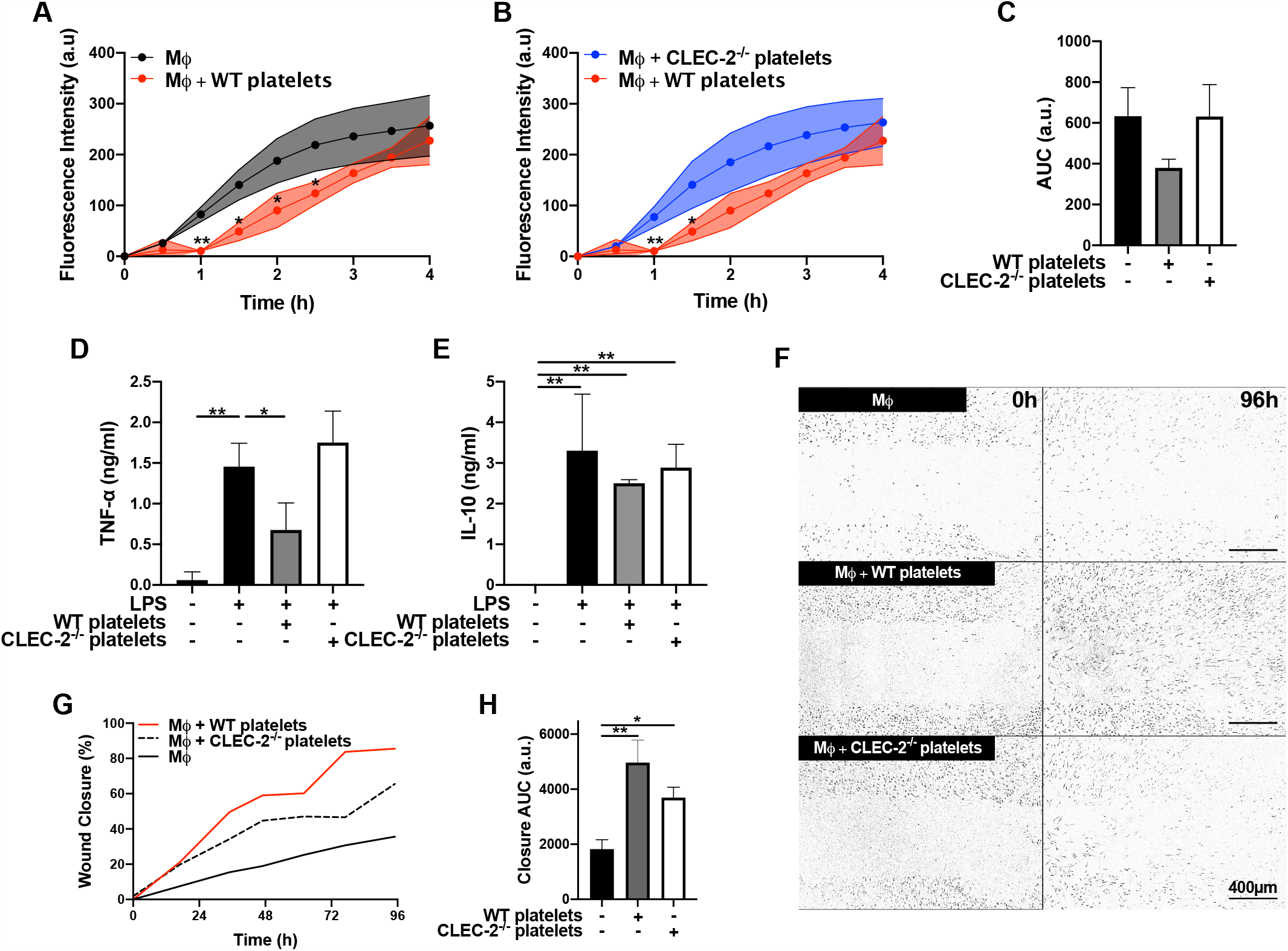
Platelet CLEC-2 delays inflammatory BMDM phagocytic capacity and reduces TNF-α Secretion. **(A-C)** pH sensitive Alexa Fluor-488 conjugated *Escherichia Coli* bioparticles (3×10^6^ beads/condition) were added to LPS-treated BMDMs (Mϕ) in the absence or presence of **(B)** WT platelets or **(C)** CLEC-2-deficient platelets (100 platelets: 1 Mϕ) for 4h. **(A, B)** Phagocytosis was visualised and quantified by time lapse-imaging using an Incucyte Live-cell analysis system. **(C)** Phagocytosis profiles were quantified at 4h by detecting fluorescence/mm^3^ using area under the curve (AUC; a.u. = arbitrary units). **(D)** TNF-α and **(E)** IL-10 secretion from BMDMs or Mϕ cultured with WT or CLEC-2-deficient platelets was quantified in the supernatant by ELISA (n=4). **(F-H)** Scratch wound migration of Mϕ was monitored every 2h for 96h using an Incucyte ZOOM system. Following wound scratch, **(F)** WT or CLEC-2-deficient platelets (100 platelets:1 Mϕ) were added to Mϕ. **(G)** Wound closure was quantified as percentage of closure using ImageJ. **(H)** Total wound closure was quantified by area under the curve (AUC) at 96h (n=3). The statistical significance between 2 groups was analyzed using a student’s paired t-test and the statistical difference between multiple groups using one-way ANOVA with Tukey’s multiple comparisons test. \**p* < 0.05 \*\**p* < 0.01.

Alongside phagocytic activity, we investigated how platelet CLEC-2 regulates BMDM migration *in vitro* using a scratch wound healing and Boyden chamber assays. A wound was made in a monolayer of inflammatory BMDMs before addition of WT or CLEC-2-deficient platelets (**Fig. 2 F**). LPS-treated macrophages displayed a 40% wound closure over 96h, which was increased to 80% when cultured with WT platelets (**Fig. 2 F-H**). Wound closure was also accelerated in the presence of CLEC-2-deficient platelets, although the maximal coverage did not exceed 60%.

Together these data suggest that platelet CLEC-2 binding to inflammatory macrophages alters cytoskeleton rearrangement, delays phagocytosis and increases macrophage migration, whilst reducing macrophage TNF-α secretion.

### Crosslinking podoplanin by rCLEC-2-Fc is sufficient to promote macrophage migration and modulate their inflammatory phenotype

Platelet-mediated alteration in inflammatory macrophage migration and function is dependent on CLEC-2. In order to assess whether CLEC-2-dependent platelet activation and secretion, or crosslinking podoplanin is responsible for these changes, we used recombinant dimeric CLEC-2 (rCLEC-2-Fc) (Bourne et al., 2020) on inflammatory macrophages and compared the effect to an IgG control. Addition of rCLEC-2-Fc to LPS-stimulated RAW264.7 cells (**Fig. 3 A)** or LPS-treated BMDMs for 1h increased podoplanin expression similar to WT platelets **(Fig. 3 B)**. Crosslinking podoplanin with rCLEC-2-Fc is associated with a decrease in TNF-α secretion (**Fig. 3 C**), suggesting that crosslinking podoplanin by CLEC-2 is responsible, rather than platelet activation and secretion, for the immunomodulatory effect of CLEC-2.

**Figure 3:**
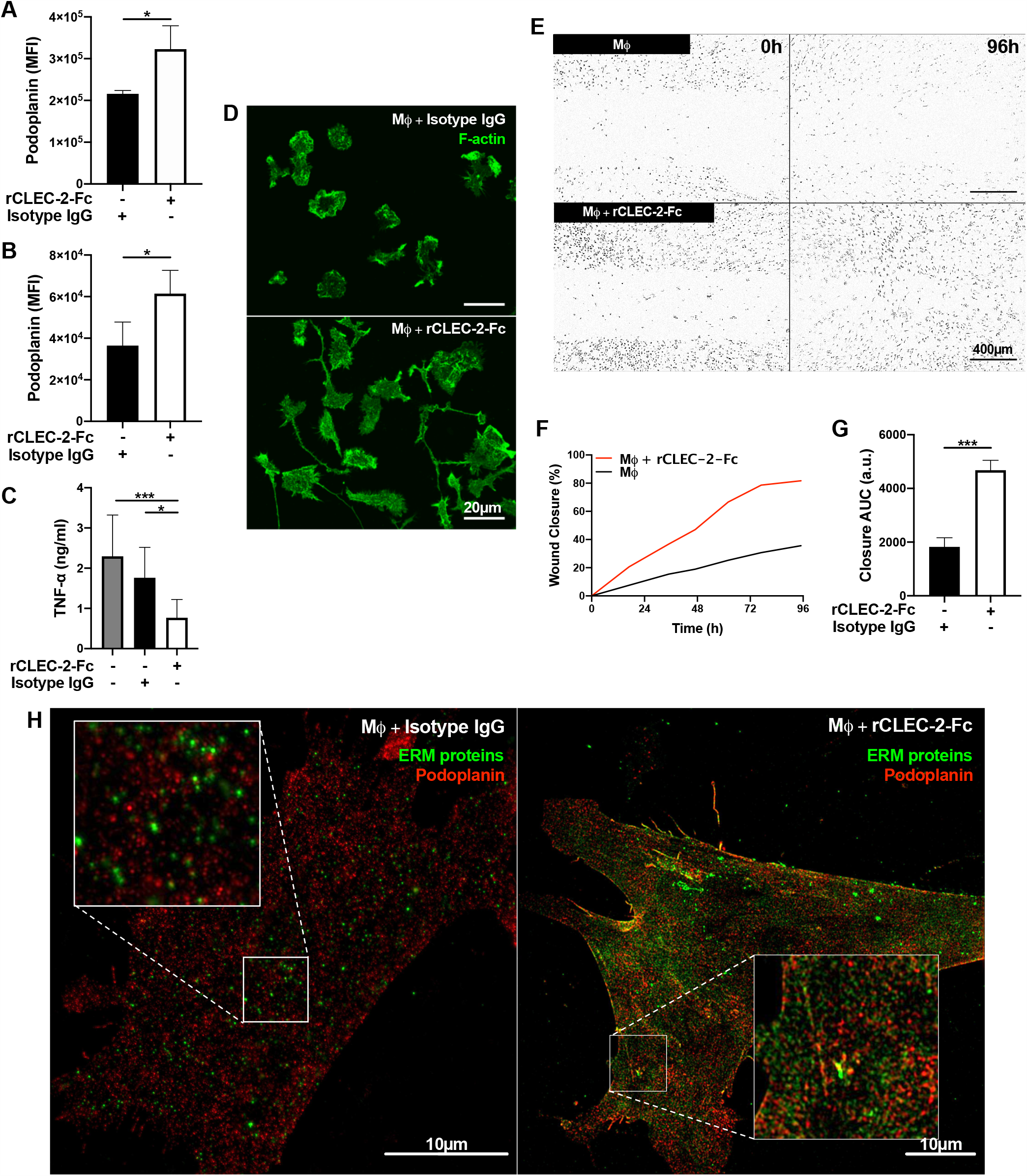
rCLEC-2-Fc upregulates podoplanin expression, and promotes macrophage spreading and migration *in vitro*. 24h LPS-stimulated (1µg/ml) **(A)** RAW264.7 cells (n=3) or **(B)** BMDMs (Mϕ; n=4), were washed and incubated with recombinant CLEC-2-Fc (rCLEC-2-Fc; 10µg/ml) or IgG isotype control (10µg/ml) for 1h. **(A, B)** Surface podoplanin expression was detected by flow cytometry using a conjugated anti-podoplanin antibody. **(C)** Mϕ were washed, and cultured with rCLEC-2-Fc (10µg/ml) or IgG isotype control (10µg/ml) before TNF-α secretion was quantified in the media supernatant by ELISA (n=4). **(D)** LifeAct-GFP-derived Mϕ were spread on collagen in the presence of rCLEC-2-Fc or IgG isotype control (10µg/ml) for 2h and measured with immunofluorescence by confocal microscopy. **(E-G)** Scratch wound migration of Mϕ was monitored every 2h for a total of 96h using an Incucyte ZOOM system. Following wound scratch, **(E)** rCLEC-2-Fc (10µg/ml) was added to Mϕ. **(F)** Wound closure was quantified as percentage of closure compared to initial scratch size using ImageJ. **(G)** Total wound closure was quantified using area under the curve (AUC) at 96h (a.u.= arbitrary units; n=3). **(H)** Mϕ were cultured on collagen in the presence of rCLEC-2-Fc or IgG isotype control (10µg/ml) for 2h. Podoplanin (red) and ERM protein (green) localisation was visualised using immunofluorescence by 3D-structured illumination microscopy (3D-SIM). The statistical significance between 2 groups was analyzed using a student’s paired t-test and the statistical difference between multiple groups using one-way ANOVA with Tukey’s multiple comparisons test. \**p* < 0.05 \*\*\**p* < 0.001.

We further investigated the effect of rCLEC-2-Fc on macrophage spreading. LPS-stimulated BMDMs from LifeAct-GFP mice were spread on collagen for 2h in the presence of rCLEC-2-Fc or isotype IgG control (**Fig. 3 D**). rCLEC-2-Fc induced spreading and elongation of inflammatory BMDMs along collagen fibres compared to isotype control, suggesting that rCLEC-2-Fc is sufficient to induce cytoskeleton rearrangement. Increased podoplanin expression is associated with accelerated wound closure (**Fig. 3 E-G**). As podoplanin is known to bind to the ERM proteins, we investigated whether rCLEC-2-Fc alters the interaction and distribution of the ERM proteins in inflammatory macrophages using 3D-SIM microscopy. In LPS-treated BMDMs, podoplanin is expressed on macrophages, with limited colocalization with ERM proteins. Addition of rCLEC-2-Fc increased the expression of the ERM proteins and the colocalization of podoplanin with ERM proteins **(Fig. 3 H)** and localisation at the filopodia.

Altogether, our results show that rCLEC-2-Fc is sufficient to induce macrophage spreading by increasing the expression of podoplanin and CD44, interaction with ERM proteins, and localization at the pseudopods of the inflammatory macrophages.

### rCLEC-2-Fc decreases inflammatory macrophage numbers in the peritoneum during endotoxemia

Using different peritonitis models, podoplanin expression was upregulated on inflammatory peritoneal macrophages following zymosan (Hou et al., 2010), LPS injection or caecal ligation and puncture (Rayes et al., 2017). We assessed the relevance of crosslinking podoplanin on macrophage migration using rCLEC-2-Fc during ongoing peritonitis induced by LPS. WT mice were injected with LPS (10mg/kg) or saline for 22h to allow inflammatory macrophage recruitment to the inflamed peritoneum. 18h post-LPS, rCLEC-2-Fc or IgG isotype control were intraperitoneally injected for an additional 4h.

We compared myeloid and lymphoid immune cell populations in the spleen and peritoneum during rCLEC-2-Fc treatment following LPS challenge. The gating strategy to identify myeloid cell populations in the peritoneal lavage fluid (PLF) is shown in **Fig. 4 A**. At 22h post LPS, no significant change in the total number of cells in the peritoneum was observed (**Fig. 4 B**) in LPS-treated mice compared to control mice. LPS injection did not significantly alter CD45^+^ cell count in the PLF compared to saline-treated mice. However, a significant reduction in CD45^+^ was observed in LPS-treated mice 4h following rCLEC-2-Fc injection (**Fig. 4 C**). The change observed in rCLEC-2-Fc-treated mice is due to a significant reduction in CD11b^+^ myeloid cells (**Fig. 4 D**) and in particular F4/80^+^ macrophages (**Fig. 4 E**). The reduction in inflammatory F4/80^+^ cells was not due to increased cell apoptosis or death as measured by AnnexinV and Sytox staining (**Fig. S4 A**). F4/80^+^ macrophages detected in the peritoneum express podoplanin, and the expression was not significantly altered by rCLEC-2-Fc treatment (**Fig. 4 F**). LPS injection increases platelet-macrophage complexes in the PLF, but this was not altered by rCLEC-2-Fc (**Fig. S4 B**). However, a significant increase in F4/80^+^ CLEC-2^+^ macrophages is observed following rCLEC-2-Fc treatment (**Fig. 4 G and H**) confirming the binding of rCLEC-2-Fc to podoplanin and macrophages without alteration in platelet-macrophage interactions. The reduction in CD11b^+^ cells in the peritoneum upon rCLEC-2-Fc treatment was not a result of a modification in neutrophil or monocyte cell count in the PLF (**Fig. S4 C and D**). No significant change in CD4^+^ or CD8^+^ T cells or CD19^+^ B cells was observed in the peritoneum post-LPS-challenge or rCLEC-2-Fc treatment (**Fig. S4 E-G)**. Similarly, no alteration in CD45^+^CD11b^-^CD11c^+^ dendritic cells or red blood cells (Ter119^+^) was observed (**Fig. S4 H and I**), suggesting the rCLEC-2-Fc does not induce bleeding in the peritoneum.

**Figure 4:**
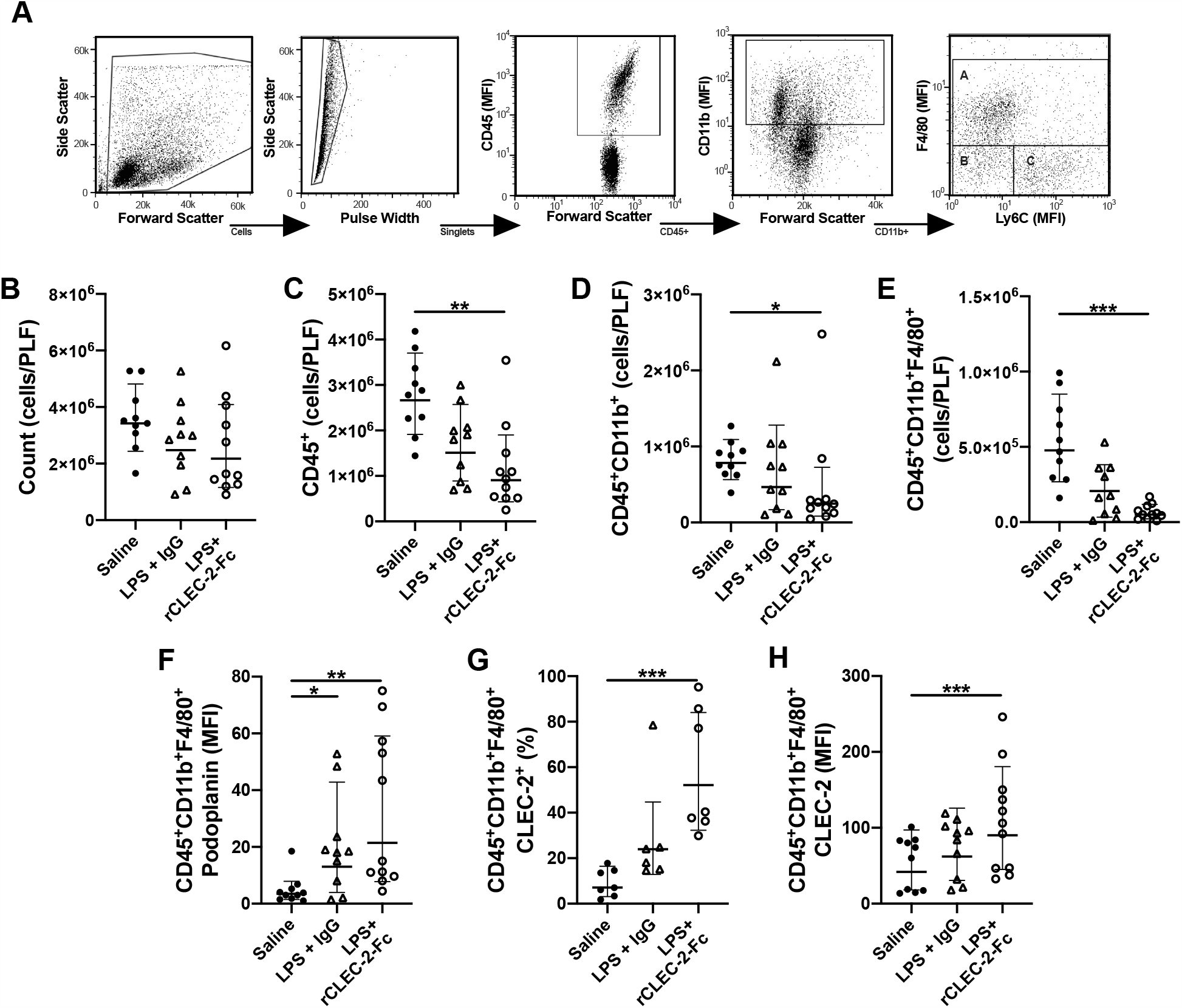
rCLEC-2-Fc decreases macrophage number in the peritoneum following endotoxemia. WT mice were intraperitoneally injected with LPS (10mg/kg) for 18h followed by rCLEC-2-Fc or IgG isotype control (100µg/mouse) for and additional 4h (n=11). Immune cell and platelet infiltration in the peritoneal lavage (PLF) were measured using flow cytometry. **(A)** Gating strategy to identify A: macrophages (F4/80^+^), B: monocytes (Ly6C^+^) and C: neutrophils (F4/80^-^Ly6C^-^). **(B)** Total number of cells from the peritoneal lavage, **(C)** leukocytes (CD45^+^), **(D)** myeloid cells (CD45^+^CD11b^+^) and **(E)** macrophages (CD45^+^CD11b^+^F4/80^+^) were measured in the PLF. **(F)** MFI of podoplanin expressed on the surface of macrophages. **(G)** Percentage of CLEC-2-positive macrophages and (**H)** the MFI of CLEC-2 expression on macrophages. The statistical significance was analyzed using a Kruskal-Wallis multiple comparisons test. \**p* < 0.05 \*\**p* < 0.01 \*\*\**p* < 0.001.

These results show that rCLEC-2-Fc selectively alters peritoneal podoplanin-positive inflammatory macrophage retention in the inflamed peritoneum.

### rCLEC-2-Fc alters inflammatory cytokine and chemokine secretion in the inflamed peritoneum

We assessed whether the change in macrophage numbers in the inflamed peritoneum is due to an alteration in cytokine and chemokine release. Similar to our *in vitro* observations, rCLEC-2-Fc decreases TNF-α secretion in the PLF of LPS-treated mice (**Fig. 5 A**) and increased the levels of the anti-inflammatory cytokine IL-10 (**Fig. 5 B**) and chemokines CCL2, CCL5 and CXCL1 (**Fig. 5 C-E**). The level of CCL21, a soluble ligand for podoplanin, was not significantly altered in rCLEC-2-Fc-treated mice (**Fig. 5 F**). The migration of macrophages was not associated with increased vascular permeability, as measured by angiopoietin-2 secretion (**Fig. 5 G**). There was no change in MMP-9, CXCL2, CCL4, C5a or IL-6 secretion following rCLEC-2-Fc treatment (**Fig. 5 H-L)**.

**Figure 5:**
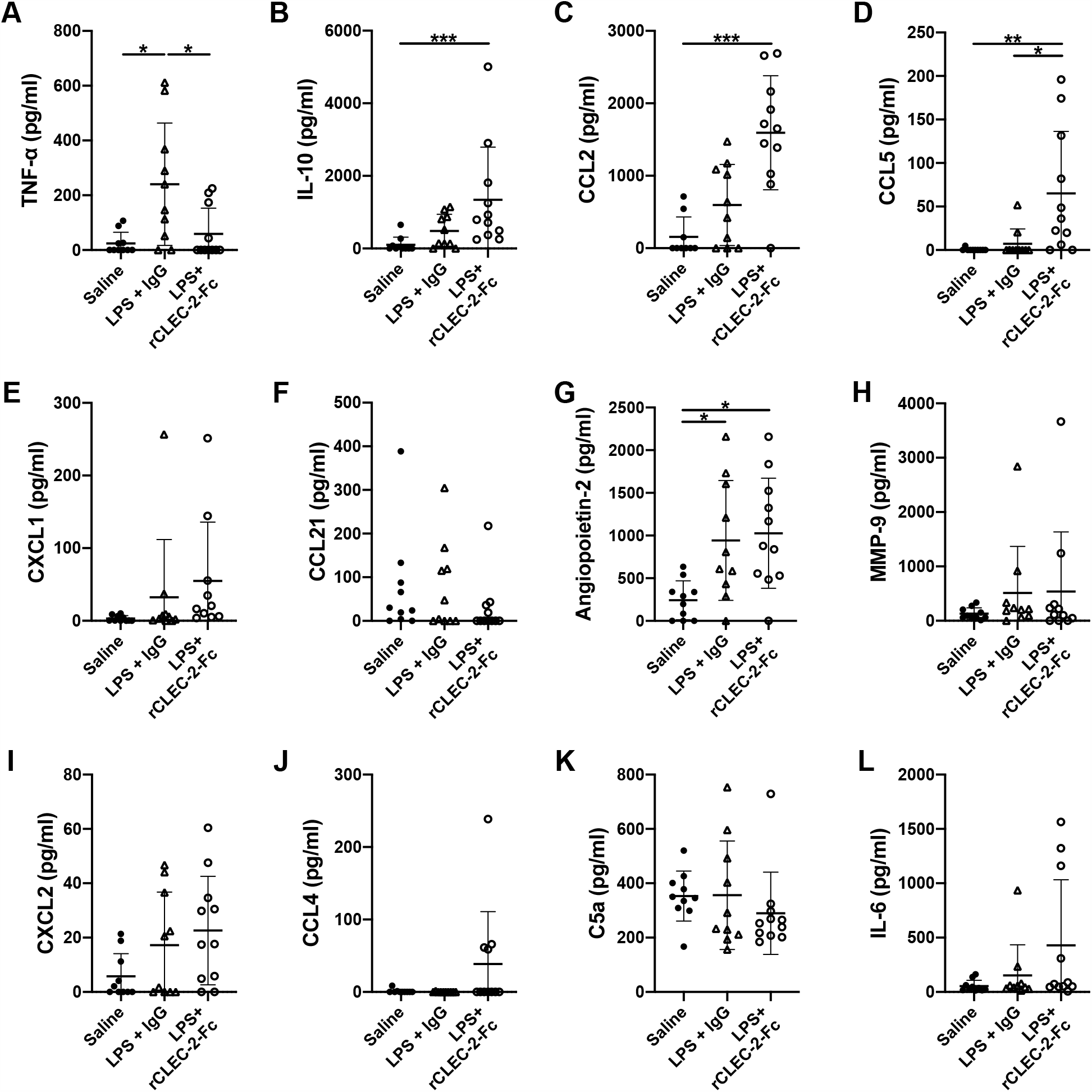
rCLEC-2-Fc alters the cytokine and chemokine profile in the peritoneum during endotoxemia and *in vitro*. WT mice were intraperitoneally injected with LPS (10mg/kg) for 18h followed by rCLEC-2-Fc or IgG isotype control (100µg/mouse) for and additional 4h (n=11). **(A-F)** Peritoneal lavage fluid (PLF) was analysed for secretion of **(A)** TNF-α, **(B)** IL-10, **(C)** CCL2, **(D)** CCL5, **(E)** CXCL1, **(F)** CCL21, **(G)** Angiopoeitin-2, **(H)** MMP-9, **(I)** CXCL2, **(J)** CCL4, **(K)** C5a and **(L)** IL-6 by ELISA. The statistical significance was analyzed using a Kruskal-Wallis multiple comparisons test for *in vivo*. \**p* < 0.05 \*\**p* < 0.01 \*\*\**p* < 0.001.

Together, our results show that injection of rCLEC-2-Fc alters cytokine and chemokine production and reduces inflammatory macrophage retention in the inflamed peritoneum.

### rCLEC-2-Fc increases podoplanin expression on F4/80^+^ macrophage in the spleen

The decrease in macrophage count in the peritoneum could be explained by increased cell death, reduced adhesion or emigration to secondary sites. Since rCLEC-2-Fc did not increase macrophage death, we first investigated the macrophage infiltration in the draining lymphoid organs. Following LPS, no significant increase in spleen weight, or CD45^+^ cell count post-LPS-challenge was observed (**Fig. 6 A and B**). However, LPS injection increases the total number of myeloid CD11b^+^ cells in the spleen compared to saline-treated mice (**Fig. 6 C**), which was abrogated in rCLEC-2-Fc-treated mice. rCLEC-2-Fc significantly decreased splenic F4/80^+^ cells numbers compared to LPS-treated mice (**Fig. 6 D**). Monocyte Ly6C^+^ and neutrophils Ly6G^+^ counts were not altered (**Fig. S5 A and B**). Similar to the inflamed peritoneum, the decrease in F4/80^+^ cells is not due to increased apoptosis or death, or alteration of macrophage-platelet complexes following rCLEC-2-Fc treatment (**Fig. S5 C and D**). Podoplanin levels on splenic macrophages in LPS-challenged mice was further increased by rCLEC-2-Fc (**Fig. 6 E**). However, the total MFI for CLEC-2 on macrophages was unchanged (**Fig. 6 F and G**). Whilst the percentage of CD41-positive macrophages is unaltered with rCLEC-2-Fc treatment, immunofluorescent imaging shows a decrease in platelet-positive macrophages in the spleen compared to IgG control (**Fig 6 H**). CD4^+^, CD8^+^, CD19^+^ and CD11c^+^ cell numbers in the spleen were not altered by rCLEC-2-Fc treatment following LPS compared to IgG treated mice (**Fig. S5 E-H)**.

**Figure 6:**
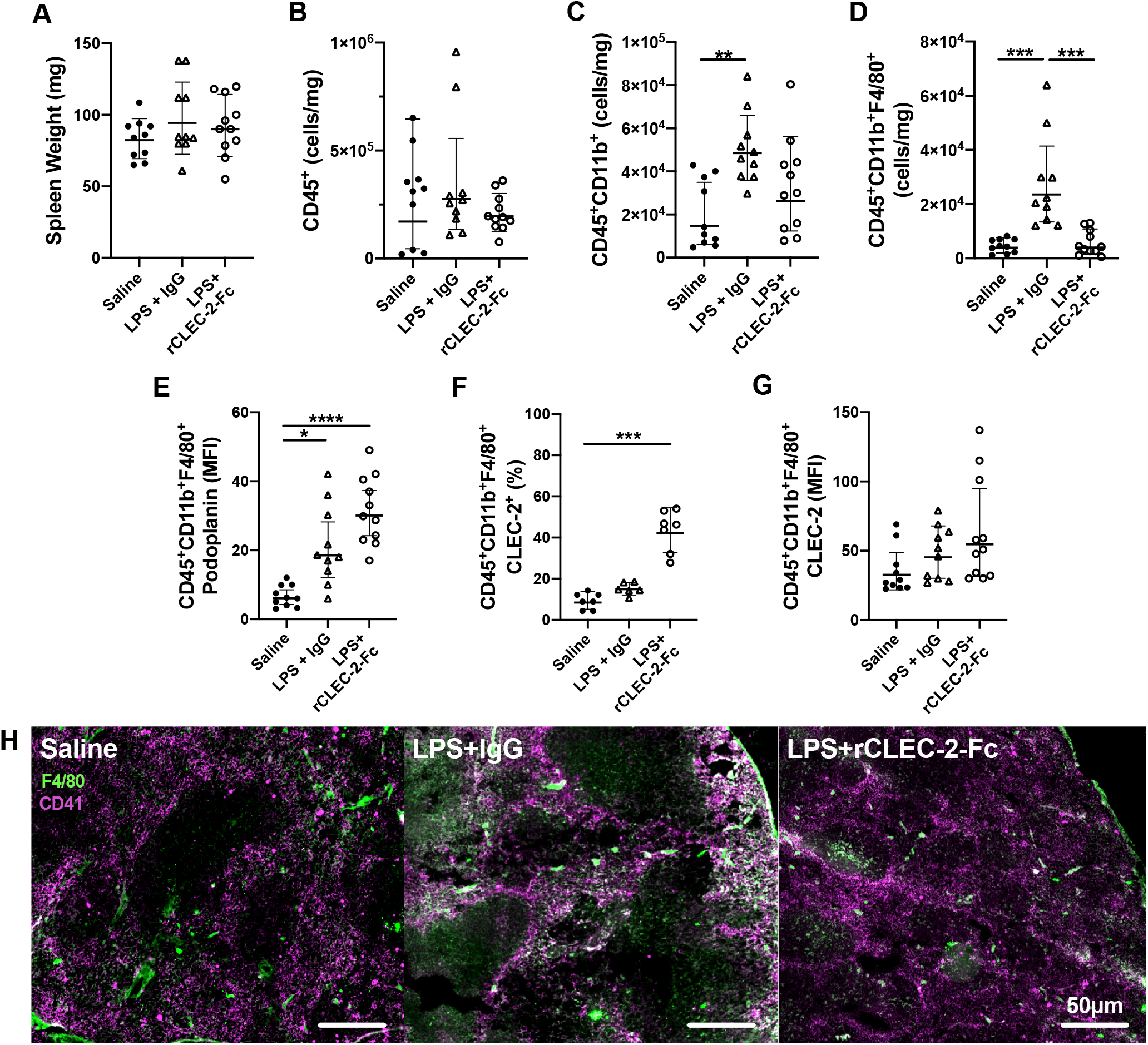
rCLEC-2-Fc does not increase macrophage infiltration into the spleen during endotoxemia. WT mice were intraperitoneally injected with LPS (10mg/kg) for 18h followed by rCLEC-2-Fc or IgG isotype control (100µg/mouse) for and additional 4h (n=11). Total splenocytes were collected, red blood cells lysed and immune cell populations detected by flow cytometry. Cell number was normalised to **(A)** spleen weight. **(B)** Total count of immune cells (CD45^+^), **(C)** myeloid cells (CD45^+^CD11b^+^) and **(D)** macrophages (CD45^+^CD11b^+^F4/80^+^), **(E)** MFI of podoplanin expressed on the surface of macrophages, **(F)** CLEC-2-positive macrophages (%) and **(G)** MFI of CLEC-2 expression on macrophages were assessed using flow cytometry. **(H)** Immunofluorescent staining of platelets (CD41; purple) and macrophages (F4/80; green) in frozen spleen sections. The statistical significance was analyzed using a Kruskal-Wallis multiple comparisons test. \**p* < 0.05 \*\**p* < 0.01 \*\*\**p* < 0.001.

Our results show that the decrease in peritoneal macrophages following rCLEC-2-Fc is not due to their emigration to the spleen.

### rCLEC-2-Fc promotes peritoneal podoplanin^+^ macrophage migration to mesenteric lymph nodes during endotoxemia

As rCLEC-2-Fc did not induce F4/80^+^ migration to the spleen, we next investigated the possible emigration of inflammatory peritoneal macrophages to the draining lymph nodes. We assessed the F4/80^+^ population in the mesenteric lymph nodes (MLN) by flow cytometry and immunofluorescence. A significant increase in myeloid cells was observed in rCLEC-2-Fc treated mice compared to IgG control (**Fig. 7 A**). We observed a significant increase in the percentage of macrophages in the MLN upon rCLEC-2-Fc treatment compared to IgG isotype control (**Fig. 7 B**). Interestingly, a similar percentage of CLEC-2-positive F4/80^+^ cells was observed in both IgG and rCLEC-2-Fc treatment, which may be explained by the presence of macrophage-platelet complexes in the MLN (**Fig. 7 C**). However, the difference in CLEC-2 MFI between isotype and rCLEC-2-Fc-treated mice further confirm that rCLEC-2-Fc is able to bind to platelet-macrophage complexes without alteration in complex formation (**Fig. 7 D**). The increase in CLEC-2 density on F4/80+ cells positively correlates with F4/80+ frequency in the MLN (r^2^=0.67, p=0.0011; **Fig. 7 E**).

**Figure 7:**
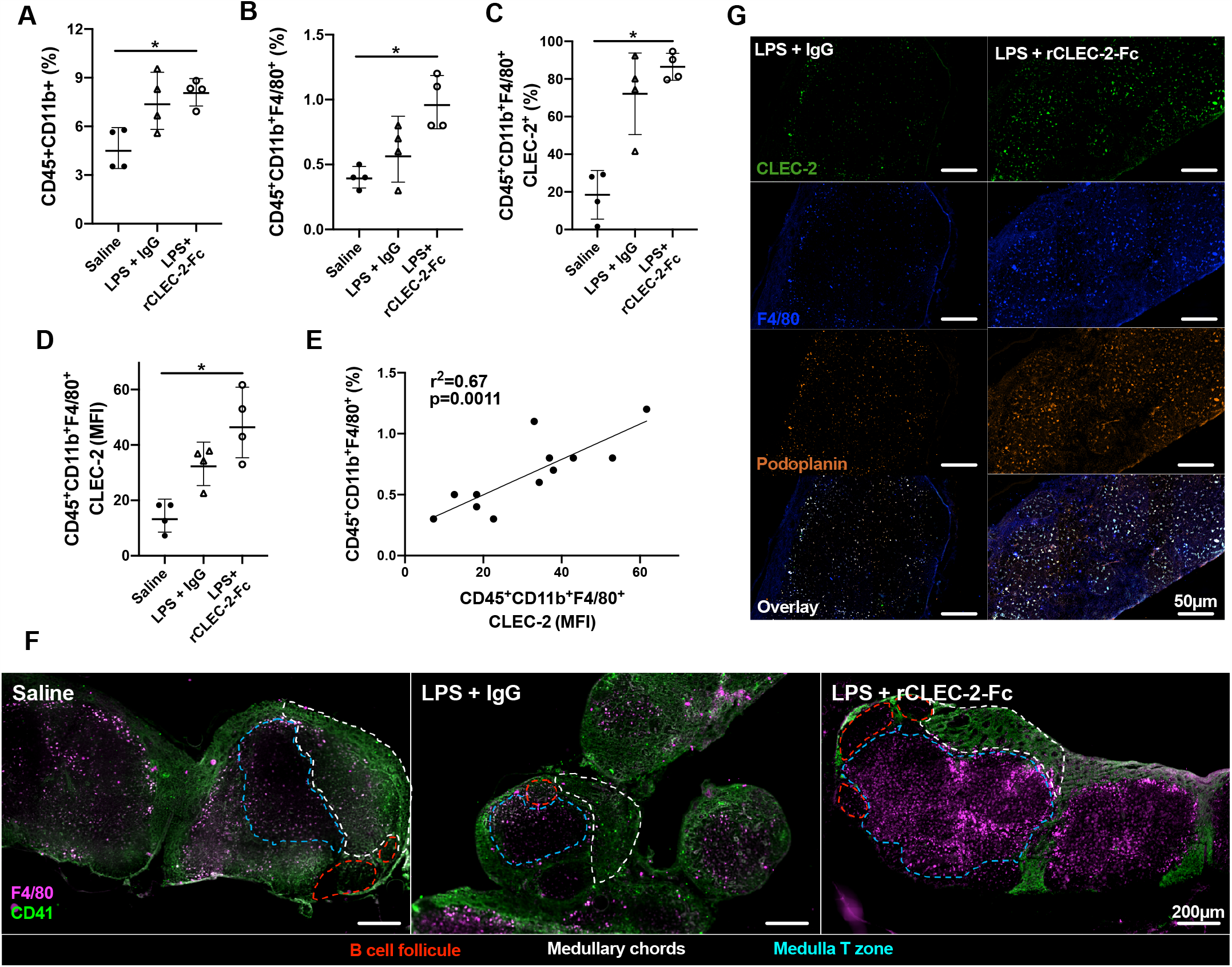
rCLEC-2-Fc increases peritoneal macrophage migration to mesenteric lymph nodes during endotoxemia. WT mice were intraperitoneally injected with LPS (10mg/kg) for 18h followed by rCLEC-2-Fc or IgG isotype control (100µg/mouse) for an additional 4h (n=4). **(A-E)** Mesenteric lymph node (MLN) cells were collected and immune cell population detected by flow cytometry. **(A)** Percentage of myeloid cells (CD45^+^CD11b^+^) and **(B)** macrophages (CD45^+^CD11b^+^F4/80^+^). **(C)** CLEC-2-positive macrophages and **(D)** the MFI of CLEC-2 expression on macrophages in MLN. **(E)** Correlation between the percentage of MLN macrophages and the MFI of CLEC-2 expression on macrophages by Simple Linear Regression. **(F)** Immunofluorescent staining of macrophages (F4/80; purple) and platelets (CD41; green) in frozen MLN sections. **(G)** Immunofluorescent staining of macrophages (F4/80; blue), CLEC-2 (green) and podoplanin (orange) in frozen MLN sections. The statistical significance was analyzed using a Kruskal-Wallis multiple comparisons test. \**p* < 0.05.

These results show that rCLEC-2-Fc binding to podoplanin^+^ peritoneal macrophages promotes their emigration from the inflamed peritoneum to the MLN.

### rCLEC-2-Fc-driven macrophage migration to mesenteric lymph nodes prime T cells

In order to assess whether emigrated macrophages prime T cells in the draining lymph nodes, we first assessed the location of these macrophages. MLNs were removed 22h post-LPS challenge and macrophages (F4/80^+^) and platelets (CD41^+^) localisation in the lymph nodes was assessed using immunofluorescence (**Fig. 7 F**). In unchallenged lymph nodes, platelets are localised in the medulla and around the medullary chords. Following LPS challenge, there is no significant staining for F4/80^+^ cells. However, following rCLEC-2-Fc treatment, a significant influx of F4/80^+^ cells is observed in the draining lymph nodes with the presence of these cells in the medulla, in close contact with T cells. We did not detect macrophages bound to platelets in this zone, however macrophages localised in the lymph nodes post-rCLEC-2-Fc are seen to be CLEC-2-and podoplanin-positive compared to IgG control (**Fig. 7 G**).

Lymphatic endothelial cells (LECs) are well reported to constituently secrete the podoplanin ligand, CCL21 (Farnsworth et al., 2019), which we investigated in the context of inflammation. CCL21 is observed co-localised to LECs in MLN, maintaining levels of secretion upon rCLEC-2-Fc treatment (**Fig. 8 A**). Alongside podoplanin, CCL21 is a ligand for receptors CCR4 and CCR7 (Van Raemdonck et al., 2020). rCLEC-2-Fc did not alter the surface expression of CCR4 or CCR7 in BMDMs (**Fig. 8 B and C**). In order to assess whether the increase in podoplanin expression by rCLEC-2-Fc alters the migration of inflammatory macrophages towards the podoplanin soluble ligand, CCL21, we assessed the effect of rCLEC-2-Fc on migration of inflammatory BMDMs using the Boyden chamber assay. Addition of rCLEC-2-Fc to LPS-treated BMDMs induced a significant increase in the number of macrophages that had migrated towards CCL21 (**Fig. 8 D**), consistent with a role for the chemokine in driving macrophage emigration from the peritoneum and into MLN.

**Figure 8:**
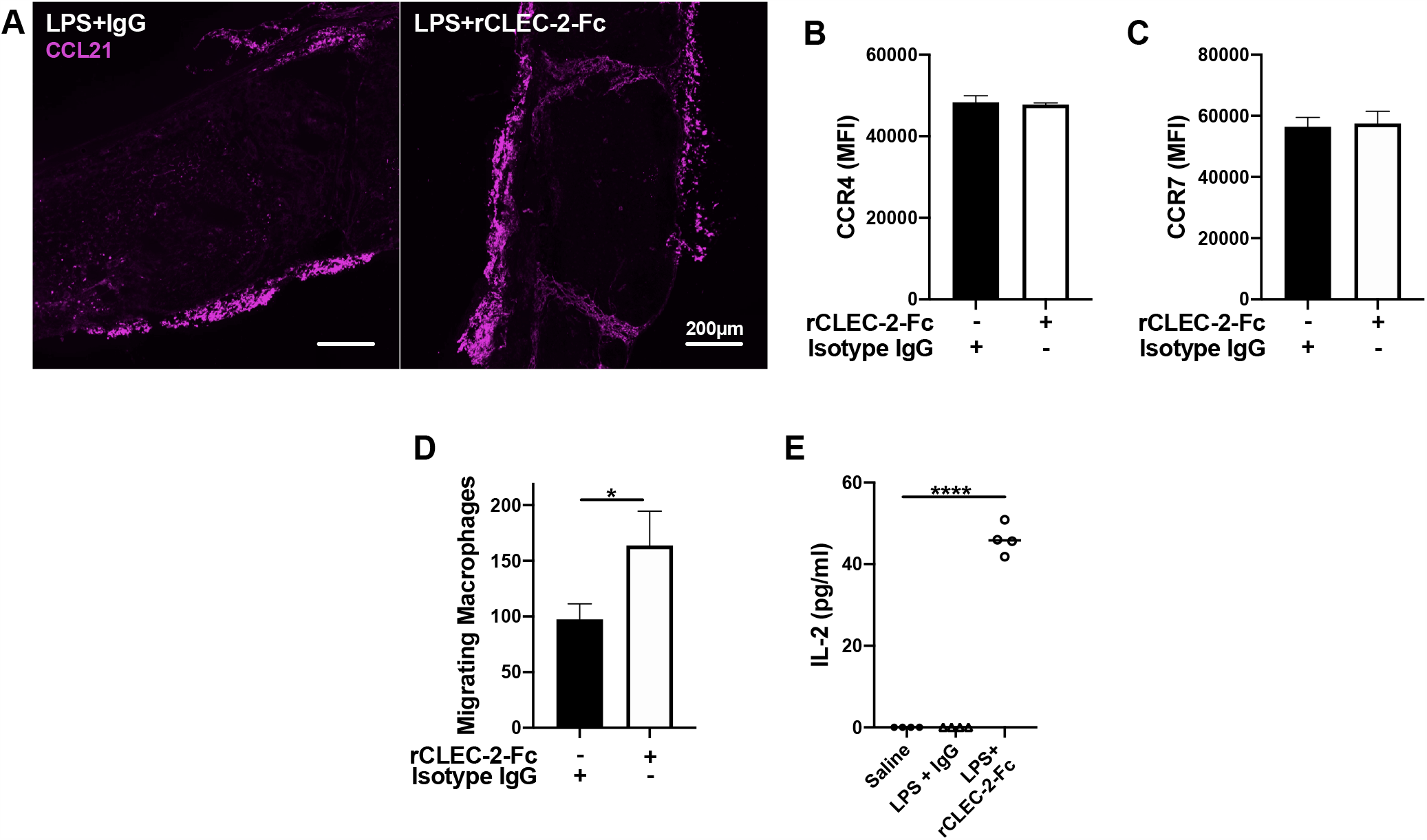
rCLEC-2-Fc increases inflammatory macrophage migration towards CCL21 and primes T cells in mesenteric lymph nodes. WT mice were intraperitoneally injected with LPS (10mg/kg) for 18h followed by rCLEC-2-Fc or IgG isotype control (100µg/mouse) for an additional 4h (n=4). **(A)** Immunofluorescent staining of CCL21 (red) in frozen MLN taken from LPS-challenged mice. **(B, C)** 24h LPS-stimulated (1µg/ml) BMDMs (Mϕ) were treated with rCLEC-2-Fc (10µg/ml) or IgG isotype control (10µg/ml) for 4h. Surface expression of **(B)** CCR4 and **(C)** CCR7 median fluorescence intensity (MFI) was detected by flow cytometry using conjugated anti-CCR4 or anti-CCR7 antibodies, respectively (n=4). **(D)** Migration of inflammatory BMDMs co-cultured with rCLEC-2-Fc or IgG Isotype control (10µg/ml) toward CCL21 (30ng/ml) was analysed using a Boyden chamber assay for 4h (n=3). **(E)** IL-2 secretion from MLN cells isolated from different mice cultured *in vitro* with LPS (10ng/ml) for 16h was quantified by ELISA.The statistical significance was analyzed using a Kruskal-Wallis multiple comparisons test. \**p* < 0.05 \*\*\*\**p* < 0.0001.

We further investigated whether infiltrated inflammatory macrophages are able to prime T cells. To test this hypothesis homogenised lymph nodes from saline- and LPS-treated mice were re-stimulated *ex vivo* with low-dose LPS (10ng/ml) for 24h, and IL-2 secretion was measured in supernatants by ELISA (**Fig. 8 E**). Low-dose LPS did not induce IL-2 secretion in the supernatant of MLN cells from saline- or LPS-treated mice. In contrast, addition of low-dose LPS to MLN cells isolated from mice treated with rCLEC-2-Fc induces IL-2 secretion in the supernatant. This is consistent with a pathway in which rCLEC-2-Fc-induces macrophage emigration to MLN, leading to the priming of T cells.

Altogether, these results suggest that rCLEC-2-Fc binding to peritoneal podoplanin-positive inflammatory macrophages promotes the interaction, and expression of podoplanin’s soluble, membrane, and intracellular ligands. This is associated with increased migration to mesenteric lymph nodes to prime T cells.

## Discussion

In this study, we report that crosslinking podoplanin using recombinant CLEC-2 promotes inflammatory podoplanin-positive macrophage migration during ongoing endotoxemia, and accelerates their removal from the site of inflammation to draining lymph nodes as well as priming of T cells. The interaction of CLEC-2 with podoplanin upregulated on inflammatory macrophages promotes their mobility through (i) the dephosphorylation of podoplanin intracellular serine residues, (ii) upregulation and rearrangement of podoplanin and CD44 expression on macrophage membrane extensions, (iii) reorganisation of actin cytoskeleton and ERM protein family distribution on cell protrusion and (iv) inflammatory macrophage interaction with CCL21 secreted by lymphatic endothelial cells (LECs). In parallel, CLEC-2 crosslinks podoplanin and reduces TNF-*α* secretion from inflammatory macrophages and delays their phagocytic activity. We propose that during acute, ongoing inflammation, CLEC-2-podoplanin crosslinking plays an anti-inflammatory role by reducing the inflammatory phenotype of macrophages. This interaction facilitates the removal of macrophages from the sites of inflammation, and hence dampens the inflammatory environment.

Using LPS-, thioglycollate- or CLP-induced peritonitis, we and other have shown a significant upregulation of podoplanin on inflammatory macrophages (Hou et al., 2010, Kerrigan et al., 2012, Rayes et al., 2017). Podoplanin interacts with many ligands such as CLEC-2, CD44 (Martín-Villar et al., 2010) and CD9, soluble proteins such as CCL21 and galectin-8 and intracellular proteins such as ERM (Martín-Villar et al., 2006). Podoplanin interaction with CLEC-2 leads to platelet activation and thrombosis. In mouse models of systemic *Salmonella Typhimurium* infection and deep vein thrombosis, upregulation of podoplanin and exposure at the site of vascular breaches triggers CLEC-2-dependent thrombus formation (Hitchcock et al., 2015, Payne et al., 2017). However, little is known on the functional relevance of this interaction on macrophage activation and mobility. Here we show that the interaction of CLEC-2 with podoplanin on inflammatory macrophages induces significant changes in macrophage migration and inflammatory phenotype.

LPS-treated RAW264.7 and BMDMs cultured with WT, but not CLEC-2-deficient platelets, increase podoplanin expression, protrusions and macrophage elongation. The effect of CLEC-2 was not dependent on platelet secretion, as recombinant dimeric CLEC-2 induces a similar phenotype, suggesting that crosslinking podoplanin is mediating the immunomodulatory effects. Deletion of CLEC-2 did not alter the binding of platelets to inflammatory macrophages. Indeed, other platelet receptors are implicated in platelet-macrophage complexes including GPIb, CD40L (Inwald et al., 2003) and P-selectin (Ye et al., 2019). GPIb-MAC-1 polarizes monocytes towards the pro-inflammatory macrophage (Carestia et al., 2019), showing that different heterotypic interactions differentially regulate macrophage function.

Platelet CLEC-2, as well as rCLEC-2-Fc, induces a rapid translocation of podoplanin from intracellular stores to the cell surface. This is associated with a loss of phosphorylation of the serine residues in the intracellular podoplanin tail, which has been demonstrated to promote fibroblast migratory activity (Krishnan et al., 2013, Krishnan et al., 2015). Classically, macrophages migrate through actin polymerisation-driven elongation of the leading edge towards a gradient, followed by integrin mediated adhesion to matrix proteins and finally actomyosin contraction and trailing edge de-adhesion (Pixley, 2012). CLEC-2 induced the reorganisation of the actin cytoskeleton and increased podoplanin interaction with ERM proteins and CD44, promoting macrophage migration. Indeed, CD44 expression is required for podoplanin-induced migration in squamous stratified epithelia (Martín-Villar et al., 2010), suggesting that CLEC-2 may increase podoplanin and CD44 expression and their association, leading to macrophage migration. Our data supports a role for rCLEC-2-Fc in inflammatory macrophage migration, as measured by wound closure assays.

In order to assess the relevance of this interaction during ongoing, acute inflammation, we injected rCLEC-2-Fc into mice following endotoxemia. Interestingly, rCLEC-2-Fc drastically decreases macrophage count in the inflamed peritoneum. The absence of macrophages from the inflamed peritoneum could be secondary to (i) macrophage local death (Gautier et al., 2013), (ii) increased macrophage adherence through integrin *α*_D_*β*_2_ and *α*_M_*β*_2_ (Mac-1) upregulation (Yakubenko et al., 2008) and/or (iii) macrophage migration to a secondary site (Bellingan et al., 1996). In our study, we show that the decrease in macrophage numbers in the inflamed peritoneum was not due to macrophage apoptosis or death. However, we observed a mass emigration of rCLEC-2-Fc bound macrophages to draining mesenteric lymph nodes (MLN) where they localise in the medulla in close proximity with T cells. The increase in macrophage emigration was not due to matrix metalloprotease 9 secretion (Gong et al., 2008), or dysregulated vascular integrity, neither by increase in other chemotactic molecules such as complement C5a levels. Indeed, CLEC-2 was previously shown to regulate the inflammatory response by increasing the secretion of complement regulators from platelets (Xie et al., 2020). We now show that CCL21, constituently secreted by LECs (Farnsworth et al., 2019), is utilised by emigrating macrophages during our model of inflammation as a gradient towards the draining MLN. CCL21 has previously been described as a key regulator of dendritic cells, also expressing podoplanin, migrating to lymph nodes (Lira, 2005, Manzo et al., 2007). The rCLEC-2-Fc-induced upregulation of podoplanin on inflammatory macrophages correlates with the increase in macrophage interaction with CCL21, and hence increase cell migration toward the chemokine’s source.

*Ex vivo* treatment of MLN cells with low doses of LPS induces IL-2 secretion in cells isolated from rCLEC-2-Fc-treated mice, but not from IgG-treated mice. This data suggests T cell priming (Sojka et al., 2004), possibly by increasing macrophage infiltration and interaction with T cells. Our data supports a model by which crosslinking podoplanin with CLEC-2 accelerates the removal of inflammatory macrophage from the site of inflammation and their migration to the draining lymph nodes. Whether the macrophage infiltration is able to promote an adaptive immune response to different antigens and alter IgG response requires further elucidation.

In conclusion, we show a novel, key immunomodulatory role for CLEC-2-podoplanin interaction through inflammatory macrophage migration and removal from the site of inflammation, independent of thrombosis. CLEC-2 crosslinking podoplanin induces a series of changes leading to macrophage migration while reducing their inflammatory phenotype. An increased inflammatory macrophage accumulation is observed in inflammatory diseases including atherosclerosis, metabolic disorders and SARS-CoV-2 infection. Crosslinking podoplanin using rCLEC-2-Fc may present a novel pathway to reduce macrophage accumulation in the inflammatory environment.

## Methods and Materials

### Mice

Wild type (WT) C57BL/6 mice (12-14 weeks; males and females) were purchased from Harlan Laboratories (Oxford, UK). Platelet-specific CLEC-2-deficient mice (Haining et al., 2020) (CLEC1b^fl/fl^GPIbCre^+^), haematopoietic-specific podoplanin deficient mice (Rayes et al., 2017) (PDPN^fl/fl^Vav-iCre^+^) and LifeAct-GFP mice (Riedl et al., 2010) were used. All experiments were performed in accordance with UK law (Animal Scientific Procedures Act 1986) with approval of the local ethics committee and UK Home Office approval under PPL P2E63AE7B, PP9677279 and P0E98D513 granted to the University of Birmingham.

### Cell culture

RAW264.7 cells (Sigma Aldrich) were cultured as previously described (Kerrigan et al., 2012). Bone marrow cells were isolated from WT mice or PDPN^fl/fl^Vav-iCre^+^ (tibias and femurs) and differentiated into bone marrow-derived macrophages (BMDM) with L-929 conditioned medium for 7 days (Weischenfeldt and Porse, 2008). RAW264.7 cells and BMDMs were maintained in Dulbecco’s Modified Eagle Media (DMEM, Thermofisher) supplemented with 10% heat-inactivated foetal bovine serum (FBS), 1% penicillin-streptomycin and 2mM L-glutamine in a humidified incubator at 5% CO_2_ and 37°C.

### Platelet preparation

Mouse platelets were prepared as previously described (Kerrigan et al., 2012).

### Recombinant protein generation

rCLEC-2-Fc was house made in our laboratory. ECD region of mouse CLEC2 was cloned into mammalian expression vector pFuse-rIgG-Fc (Invitrogen). The construct was used to stable transfect HEK-293T cells with PEI method. The stable line is established by Zeocin selection and function screening. rCLEC2-fc was then purified from the cell culture supernatant by protein-A affinity chromatography, followed validations by SDS-PAGE, western blots and various functional assays.

### Protein and transcription quantification

Antibody details are listed in **Table 1**. BMDMs and RAW264.7 cells were cultured with LPS (*Escherichia Coli* 055: B5) 1µg/ml for 24h. Macrophages were scraped in ice-cold PBS-EDTA (5mM), Fc-receptors blocked (anti-CD16/CD31) in 10% mouse serum), and proteins surface expression assessed by Accuri C6 Plus flow cytometry (BD Bioscience, Oxford, UK). Cytokines and chemokines were measured in the cell media supernatant by ELISA (Peprotech). For western blotting, cells were lysed in NP40 buffer supplemented with 1%PMSF, Na_3_VO_4_, NaF and protease inhibitor cocktail (Roche). Protein was estimated by Bradford reagent (Biorad). Protein electrophoresis was performed in Tris-Glycine SDS running buffer (Biorad). Immunoprecipitation was performed in whole cell lysate using anti-podoplanin antibody (clone 8.1.1) bound to protein A-sepharose beads. For transcriptional quantification, RNA was TRIzol-isolated (Invitrogen) and cDNA transcribed using PrimeScript (Clontech, Takara). mPodoplaninFc, the extracellular podoplanin domain, was amplified from cDNA using primers mPodoHindFor (GATCAAGCTTATGTGGACCGTGCCAGTGTTG) and mPodoFcRev (GATCGGATCCACTTACCTGTCAGGGTGACTACTGGCAAGCC) and was quantified by SYBR Green 1 mastermix (Roche) qPCR using a PCR thermocycler and normalised to unstimulated control.

### Confocal microscopy

BMDMs and RAW264.7 cells were cultured with LPS (1µg/ml) for 24h before being allowed to interact with platelets in suspension for 15min. For immunofluorescence, cells were fixed in 4% paraformaldehyde (PFA), blocked (1%BSA, 5% goat serum) and stained using for nuclei using Hoescht 33342 and specific antibodies (**Table 1**). Cells were flat-mounted using Hydromount (National Diagnostics). Images were acquired using an inverted confocal microscope (Zeiss LSM 780) using a 1.49 NA 64X oil-immersion objective. Argon-ion lasers 405nm, 457-514nm and 561nm were used to excite constructs. Cells were quantified for size and circularity using ImageJ.

### LightSheet diSPIM microscopy

LPS-activated BMDMs derived from LifeAct-GFP mice were allowed to spread on collagen coated slides. Platelets from WT mice were stained with CellMask Deep Red plasma membrane stain (Invitrogen) and were allowed to interact with the BMDMs for 15 min prior to imaging. Cells were imaged in phenol red free DMEM supplemented with 2% FBS at 37°C and 5% CO_2_. Fluorescence was acquired on a Marianas LightSheet (Intelligent Imaging Innovations, Denver, CO, USA), a dual inverted Selective Plane Illumination Microscope (diSPIM) fitted with two perpendicular 0.8 NA, 40x water immersion objectives and ORCA-Flash4.0 V3 sCMOS cameras (Hamamatsu), driven by SlideBook software (Intelligent Imaging Innovations, Denver, CO, USA). Volumes were captured every minute in slice scan mode, with a step size of 0.5µm and 488nm and 640nm excitation wavelengths. Time lapse movies of maximal intensity projections were visualised using Arivis.

### 3D-Structured illumination microscopy (SIM)

BMDMs were allowed to spread on collagen-coated #1.5H glass coverslips for 2h. Cells were then fixed prewarmed 4% PFA in PEM buffer, permeabilised and blocked before immunofluorescent staining (ERM-AF488; podoplanin-AF568) and flat-mounting with VECTASHIELD antifade mounting medium (Vector Labs). SIM imaging was performed on a Nikon N-SIM-S system with Nikon Perfect Focus in 3-D SIM mode, using a Nikon 1.49x Na oil Objective, Chroma ET525/50m and ET595/50m excitation filters, and an ORCA Flash4.0 sCMOS camera (Hamamatsu). Illumination was with 488nm and 561nm lasers. Capture and subsequent reconstruction was performed in Nikon NIS Elements 5 – stack and a maximum score of less than 8 were discarded. Composite Images were visualised and adjusted for figures using ImageJ.

### Phagocytosis

LPS-treated BMDMs were incubated in the absence or presence of WT or CLEC-2-deficient platelets (100 platelets: 1 Mϕ) for 1h. pH sensitive, AlexaFluor488-conjugated, *Escherichia Coli* covered (K-12 strain) BioParticles (3⨯10^6^ beads/condition) were added for an additional 4h at 37°C. Total green fluorescence was quantified by a mask generated by Incucyte Systems on images taken using an Incucyte SX-5 Live-Cell microscope (Satorius).

### Wound healing

Confluent, LPS-treated BMDMs grown in DMEM were washed and scratched using a 96-pin 800µm-wide mechanical WoundMaker (Grada et al., 2017). They were then co-cultured in the absence or presence of WT or CLEC-2-deficient platelets (100:1) or rCLEC-2-Fc (10µg/ml) for 96h at 37°C. Cells were imaged every 2h using an Incucyte SX-5 Live-Cell microscope (Satorius). Wound closure was quantified using ImageJ, and calculated as percentage of closure at each timepoint by:

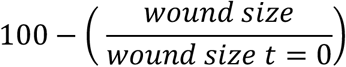

### Boyden Chamber transwell assay

For BMDM chemoattractant stimulated migration, LPS-treated BMDMs were incubated for 1h with rCLEC-2-Fc (10µg/ml) or IgG isotype (10µg/ml). Cells were scraped and seeded onto the upper layer of a 8µm pore polycarbonate inserts transwell (Corning). Migration of BMDMs towards CCL21 (30ng/ml) was assessed after 4h at 37°C. Cells were fixed, stained with Hoescht 33342 and migrated cell nuclei counted.

### LPS-induced endotoxemia in mice

LPS (*Escherichia Coli* 055:B5, Sigma) was injected intraperitoneally (IP) to age and sex matched C57BL/6 mice in 200µl saline (10mg/kg). 18h post-LPS-challenge, mice received an intraperitoneal injection of rCLEC-2-Fc or IgG isotype control (100µg/mouse) for an additional 4h. EDTA-treated blood was used to assess blood haematological parameters. Spleen and lymph nodes were homogenised and Ammonium-Chloride-Potassium (ACK) treated (5mM) and Fc-receptors blocked before antibody staining. Peritoneal cells were collected in 2ml of PBS-EDTA (10mM). Peritoneal lavage fluid (PLF), spleen and lymph node cells were measured using a CyAn ADP High-Performance Flow Cytometer.

### Immunofluorescent tissue staining

For frozen sections, organs were fixed in 4% PFA for 2h, and transferred to 20% sucrose overnight, and frozen in optimal cutting temperature (OCT) compound (Tissue-TEK, Netherlands). Organs were sectioned by cryostat at 6 microns for immunofluorescent staining. Frozen sections were acetone fixed, autofluorescence quenched with ammonium chloride (20mM), blocking (1% BSA, 5% goat serum) and staining with primary antibodies (**Table 1**), followed by secondary fluorescent antibodies. Sections were flat-mounted with VECTASHIELD antifade mounting medium (Vector Labs). Images were acquired using a Zeiss AxioScan.Z1 microscope and analysed using ZEN software.

### Data analysis

All data is presented as mean±SEM. The statistical significance between 2 groups was analyzed using a student’s paired t-test and the statistical difference between multiple groups *in* vitro using one-way ANOVA with Tukey’s multiple comparisons test. The statistical significance for *in vivo* work was determined by a Kruskal-Wallis test using Prism 8 (GraphPad Software Inc, USA). Statistical significance was represented by stars: \**p*<0.05 \*\**p*<0.01 \*\*\**p*<0.001 ****<0.0001.

## Acknowledgments

The authors would like to thank Dr Beata Grygielska for genotyping of mice, Dr Elisabeth Haining for colony maintenance, and Dr Christopher Smith for assistance throughout. This work was supported by a College-funded PhD studentship (University of Birmingham), a BHF Accelerator Award (AA/18/2/34218) and the Centre of Membrane Proteins and Receptors. JR is supported by a BHF Intermediate Basic Science Fellowship Application (FS/IBSRF/20/25039). AB is supported by BHF Senior basic Science Research Fellowship (FS/19/30/34173). SPW holds a BHF Chair (CH/03/003). Visual abstract was created using Servier Medical Art templates, licensed under Creative Commons Attribution 3.0 Unported License; https://smart.servier.com

## Contributions

JHB designed and performed research, collected, analysed and interpreted data and wrote the manuscript; NB performed experiments and contributed to data analysis; MZ, JC, EG and MC performed experiments; YD and SGT provided reagents; LVT contributed to data analysis; AB and JB contributed to data interpretation; SPW contributed to data interpretation and provided reagents; JR designed and performed research, interpreted data and wrote the manuscript. All authors reviewed and approved the manuscript.

## Data and materials availability

All data associated with this study are available in the main text or the supplementary materials.

## References

Bellingan, G. J., H. Caldwell, S. E. Howie, I. Dransfield & C. Haslett 1996. In vivo fate of the inflammatory macrophage during the resolution of inflammation: inflammatory macrophages do not die locally, but emigrate to the draining lymph nodes. J Immunol, 157, 2577–85.

Bertozzi, C. C., A. A. Schmaier, P. Mericko, P. R. Hess, Z. Zou, M. Chen, C. Y. Chen, B. Xu, M. M. Lu, D. Zhou, E. Sebzda, M. T. Santore, D. J. Merianos, M. Stadtfeld, A. W. Flake, T. Graf, R. Skoda, J. S. Maltzman, G. A. Koretzky & M. L. Kahn 2010. Platelets regulate lymphatic vascular development through CLEC-2-SLP-76 signaling. Blood, 116, 661–70.

Bourne, J. H., M. Colicchia, Y. Di, E. Martin, A. Slater, L. T. Roumenina, J. D. Dimitrov, S. P. Watson & J. Rayes 2020. Heme induces human and mouse platelet activation through C-type-lectin-like receptor-2. Haematologica.

Bretscher, A., K. Edwards & R. G. Fehon 2002. ERM proteins and merlin: integrators at the cell cortex. Nat Rev Mol Cell Biol, 3, 586–99.

Carestia, A., H. A. Mena, C. M. Olexen, J. M. OrtizWilczyñski, S. Negrotto, A. E. Errasti, R. M. Gómez, C. N. Jenne, E. A. Carrera Silva & M. Schattner 2019. Platelets Promote Macrophage Polarization toward Pro-inflammatory Phenotype and Increase Survival of Septic Mice. Cell Rep, 28, 896-908.e5.

Farnsworth, R. H., T. Karnezis, S. J. Maciburko, S. N. Mueller & S. A. Stacker 2019. The Interplay Between Lymphatic Vessels and Chemokines. Front Immunol, 10, 518.

Farr, A. G., M. L. Berry, A. Kim, A. J. Nelson, M. P. Welch & A. Aruffo 1992. Characterization and cloning of a novel glycoprotein expressed by stromal cells in T-dependent areas of peripheral lymphoid tissues. J Exp Med, 176, 1477–82.

Gautier, E. L., S. Ivanov, P. Lesnik & G. J. Randolph 2013. Local apoptosis mediates clearance of macrophages from resolving inflammation in mice. Blood, 122, 2714–22.

Gong, Y., E. Hart, A. Shchurin & J. Hoover-Plow 2008. Inflammatory macrophage migration requires MMP-9 activation by plasminogen in mice. J Clin Invest, 118, 3012–24.

Grada, A., M. Otero-Vinas, F. Prieto-Castrillo, Z. Obagi & V. Falanga 2017. Research Techniques Made Simple: Analysis of Collective Cell Migration Using the Wound Healing Assay. J Invest Dermatol, 137, e11–e16.

Greenberg, S. & S. Grinstein 2002. Phagocytosis and innate immunity. Curr Opin Immunol, 14, 136–45.

Haining, E. J., K. L. Lowe, S. Wichaiyo, R. P. Kataru, Z. Nagy, D. P. Kavanagh, S. Lax, Y. Di, B. Nieswandt, B. Ho-Tin-Noé, B. J. Mehrara, Y. A. Senis, J. Rayes & S. P. Watson 2020. Lymphatic blood filling in CLEC-2-deficient mouse models. Platelets, 1–16.

Hatzioannou, A., S. Nayar, A. Gaitanis, F. Barone, C. Anagnostopoulos & P. Verginis 2016. Intratumoral accumulation of podoplanin-expressing lymph node stromal cells promote tumor growth through elimination of CD4. Oncoimmunology, 5, e1216289.

Hitchcock, J. R., C. N. Cook, S. Bobat, E. A. Ross, A. Flores-Langarica, K. L. Lowe, M. Khan, C. C. Dominguez-Medina, S. Lax, M. Carvalho-Gaspar, S. Hubscher, G. E. Rainger, M. Cobbold, C. D. Buckley, T. J. Mitchell, A. Mitchell, N. D. Jones, N. Van Rooijen, D. Kirchhofer, I. R. Henderson, D. H. Adams, S. P. Watson & A. F. Cunningham 2015. Inflammation drives thrombosis after Salmonella infection via CLEC-2 on platelets. J Clin Invest, 125, 4429–46.

Hou, T. Z., J. Bystrom, J. P. Sherlock, O. Qureshi, S. M. Parnell, G. Anderson, D. W. Gilroy & C. D. Buckley 2010. A distinct subset of podoplanin (gp38) expressing F4/80+ macrophages mediate phagocytosis and are induced following zymosan peritonitis. FEBS Lett, 584, 3955–61.

Huang, X., F. Venet, Y. L. Wang, A. Lepape, Z. Yuan, Y. Chen, R. Swan, H. Kherouf, G. Monneret, C. S. Chung & A. Ayala 2009. PD-1 expression by macrophages plays a pathologic role in altering microbial clearance and the innate inflammatory response to sepsis. Proc Natl Acad Sci U S A, 106, 6303–8.

Inoue, O., K. Hokamura, T. Shirai, M. Osada, N. Tsukiji, K. Hatakeyama, K. Umemura, Y. Asada, K. Suzuki-Inoue & Y. Ozaki 2015. Vascular Smooth Muscle Cells Stimulate Platelets and Facilitate Thrombus Formation through Platelet CLEC-2: Implications in Atherothrombosis. PLoS One, 10, e0139357.

Inwald, D. P., A. Mcdowall, M. J. Peters, R. E. Callard & N. J. Klein 2003. CD40 is constitutively expressed on platelets and provides a novel mechanism for platelet activation. Circ Res, 92, 1041–8.

Italiani, P. & D. Boraschi 2014. From Monocytes to M1/M2 Macrophages: Phenotypical vs. Functional Differentiation. Front Immunol, 5, 514.

Kaplan, M. J. & M. Radic 2012. Neutrophil extracellular traps: double-edged swords of innate immunity. J Immunol, 189, 2689–95.

Kerrigan, A. M., L. Navarro-Nuñez, E. Pyz, B. A. Finney, J. A. Willment, S. P. Watson & G. D. Brown 2012. Podoplanin-expressing inflammatory macrophages activate murine platelets via CLEC-2. J Thromb Haemost, 10, 484–6.

Krishnan, H., J. A. Ochoa-Alvarez, Y. Shen, E. Nevel, M. Lakshminarayanan, M. C. Williams, M. I. Ramirez, W. T. Miller & G. S. Goldberg 2013. Serines in the intracellular tail of podoplanin (PDPN) regulate cell motility. J Biol Chem, 288, 12215–21.

Krishnan, H., E. P. Retzbach, M. I. Ramirez, T. Liu, H. Li, W. T. Miller & G. S. Goldberg 2015. PKA and CDK5 can phosphorylate specific serines on the intracellular domain of podoplanin (PDPN) to inhibit cell motility. Exp Cell Res, 335, 115–22.

Lira, S. A. 2005. A passport into the lymph node. Nat Immunol, 6, 866–8.

Manzo, A., S. Bugatti, R. Caporali, R. Prevo, D. G. Jackson, M. Uguccioni, C. D. Buckley, C. Montecucco & C. Pitzalis 2007. CCL21 expression pattern of human secondary lymphoid organ stroma is conserved in inflammatory lesions with lymphoid neogenesis. Am J Pathol, 171, 1549–62.

Martín-Villar, E., B. Fernández-Muñoz, M. Parsons, M. M. Yurrita, D. Megías,E. Pérez-Gómez, G. E. Jones & M. Quintanilla 2010. Podoplanin associates with CD44 to promote directional cell migration. Mol Biol Cell, 21, 4387–99.

Martín-Villar, E., D. Megías, S. Castel, M. M. Yurrita, S. Vilaró & M. Quintanilla 2006. Podoplanin binds ERM proteins to activate RhoA and promote epithelial-mesenchymal transition. J Cell Sci, 119, 4541–53.

Meng, W., A. Paunel-Görgülü, S. Flohé, A. Hoffmann, I. Witte, C. Mackenzie, S. E. Baldus, J. Windolf & T. T. Lögters 2012. Depletion of neutrophil extracellular traps in vivo results in hypersusceptibility to polymicrobial sepsis in mice. Crit Care, 16, R137.

Payne, H., T. Ponomaryov, S. P. Watson & A. Brill 2017. Mice with a deficiency in CLEC-2 are protected against deep vein thrombosis. Blood, 129, 2013–2020.

Peters, A., L. A. Pitcher, J. M. Sullivan, M. Mitsdoerffer, S. E. Acton, B. Franz, K. Wucherpfennig, S. Turley, M. C. Carroll, R. A. Sobel, E. Bettelli & V. K. Kuchroo 2011. Th17 cells induce ectopic lymphoid follicles in central nervous system tissue inflammation. Immunity, 35, 986–96.

Pixley, F. J. 2012. Macrophage Migration and Its Regulation by CSF-1. Int J Cell Biol, 2012, 501962.

Quintanilla, M., L. Montero-Montero, J. Renart & E. Martín-Villar 2019. Podoplanin in Inflammation and Cancer. Int J Mol Sci, 20.

Rayes, J., J. H. Bourne, A. Brill & S. P. Watson 2020. The dual role of platelet-innate immune cell interactions in thrombo-inflammation. Res Pract Thromb Haemost, 4, 23–35.

Rayes, J., S. Lax, S. Wichaiyo, S. K. Watson, Y. Di, S. Lombard, B. Grygielska, S. W. Smith, K. Skordilis & S. P. Watson 2017. The podoplanin-CLEC-2 axis inhibits inflammation in sepsis. Nat Commun, 8, 2239.

Retzbach, E. P., S. A. Sheehan, E. M. Nevel, A. Batra, T. Phi, A. T. P. Nguyen, Y. Kato, S. Baredes, M. Fatahzadeh, A. J. Shienbaum & G. S. Goldberg 2018. Podoplanin emerges as a functionally relevant oral cancer biomarker and therapeutic target. Oral Oncol, 78, 126–136.

Riedl, J., K. C. Flynn, A. Raducanu, F. Gärtner, G. Beck, M. Bösl, F. Bradke, S. Massberg, A. Aszodi, M. Sixt & R. Wedlich-Söldner 2010. Lifeact mice for studying F-actin dynamics. Nat Methods, 7, 168–9.

Sojka, D. K., D. Bruniquel, R. H. Schwartz & N. J. Singh 2004. IL-2 secretion by CD4+ T cells in vivo is rapid, transient, and influenced by TCR-specific competition. J Immunol, 172, 6136–43.

Suzuki-Inoue, K., Y. Kato, O. Inoue, M. K. Kaneko, K. Mishima, Y. Yatomi, Y. Yamazaki, H. Narimatsu & Y. Ozaki 2007. Involvement of the snake toxin receptor CLEC-2, in podoplanin-mediated platelet activation, by cancer cells. J Biol Chem, 282, 25993–6001.

Takakubo, Y., H. Oki, Y. Naganuma, K. Saski, A. Sasaki, Y. Tamaki, Y. Suran, T. Konta & M. Takagi 2017. Distribution of Podoplanin in Synovial Tissues in Rheumatoid Arthritis Patients Using Biologic or Conventional Disease-Modifying Anti-Rheumatic Drugs. Curr Rheumatol Rev, 13, 72–78.

Van Raemdonck, K., S. Umar, K. Palasiewicz, S. Volkov, M. V. Volin, S. Arami, H. J. Chang, B. Zanotti, N. Sweiss & S. Shahrara 2020. CCL21/CCR7 signaling in macrophages promotes joint inflammation and Th17-mediated osteoclast formation in rheumatoid arthritis. Cell Mol Life Sci, 77, 1387–1399.

Weischenfeldt, J. & B. Porse 2008. Bone Marrow-Derived Macrophages (BMM): Isolation and Applications. CSH Protoc, 2008, pdb.prot5080.

Xiang, B., G. Zhang, L. Guo, X. A. Li, A. J. Morris, A. Daugherty, S. W. Whiteheart, S. S. Smyth & Z. Li 2013. Platelets protect from septic shock by inhibiting macrophage-dependent inflammation via the cyclooxygenase 1 signalling pathway. Nat Commun, 4, 2657.

Xie, Z., B. Shao, C. Hoover, M. Mcdaniel, J. Song, M. Jiang, Z. Ma, F. Yang, J. Han, X. Bai, C. Ruan & L. Xia 2020. Monocyte upregulation of podoplanin during early sepsis induces complement inhibitor release to protect liver function. JCI Insight, 5.

Yakubenko, V. P., N. Belevych, D. Mishchuk, A. Schurin, S. C. Lam & T. P. Ugarova 2008. The role of integrin alpha D beta2 (CD11d/CD18) in monocyte/macrophage migration. Exp Cell Res, 314, 2569–78.

Ye, Z., L. Zhong, S. Zhu, Y. Wang, J. Zheng, S. Wang, J. Zhang & R. Huang 2019. The P-selectin and PSGL-1 axis accelerates atherosclerosis via activation of dendritic cells by the TLR4 signaling pathway. Cell Death Dis, 10, 507.

