## Supplementary Data for "CLEC-2 promotes inflammatory peritoneal macrophage emigration to draining lymph nodes during endotoxemia"

**Video 1 and 2: Platelets induce actin remodelling to increase spreading and pseudopod formation in inflammatory BMDMs**

Bone marrow-derived macrophages (BMDMs) from LifeAct-GFP mice were incubated with LPS (1µg/ml) for 24h (Mϕ). Mϕ were mixed in suspension in the absence **(Video 1)** or presence of CellMask deep red-stained WT platelets (red) for 15min **(Video 2)** before culturing on glass. Immunofluorescent images were acquired for 1h by LightSheet diSPIM microscopy. Movies are representative of 3 independent experiments.

### Figure S1.

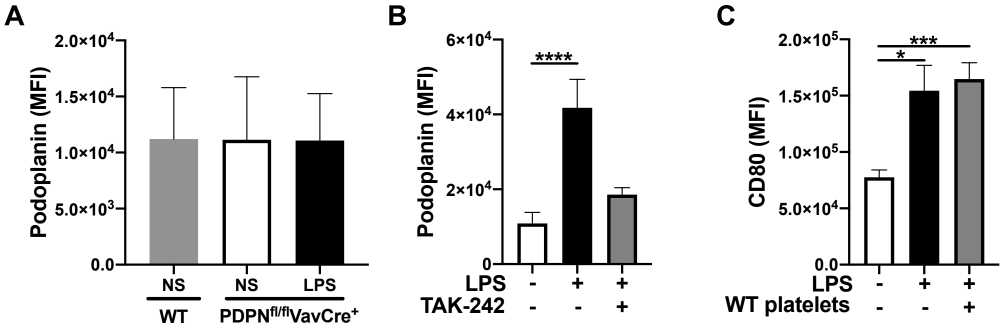


**LPS upregulates surface podoplanin expression on inflammatory BMDMs through TLR-4**

**(A)** WT or PDPN$^{fl/fl}$VAV Cre+ BMDMs were stimulated with LPS (1µg/ml) for 24h. **(B)** BMDMs were pre-treated with TAK-242 (1µM) for 1h prior to LPS stimulation (1µg/ml). **(A, B)** Podoplanin expression was compared to non-stimulated (NS) BMDMs by flow cytometry using an anti-podoplanin antibody (n=4). **(C)** BMDMs were culture in the absence or presence of LPS for 24h (1µg/ml) before washing, and co-culturing with wild type (WT) platelets (100 platelets:1BMDM). CD80 expression was acquired by flow cytometry, using an anti-CD80 antibody (n=3). The statistical significance was analyzed using a one-way ANOVA with Tukey's multiple comparisons test. *$p < 0.05$ **$p < 0.01$ ***$p < 0.001$ ****$p < 0.0001$.

#### Figure S2.

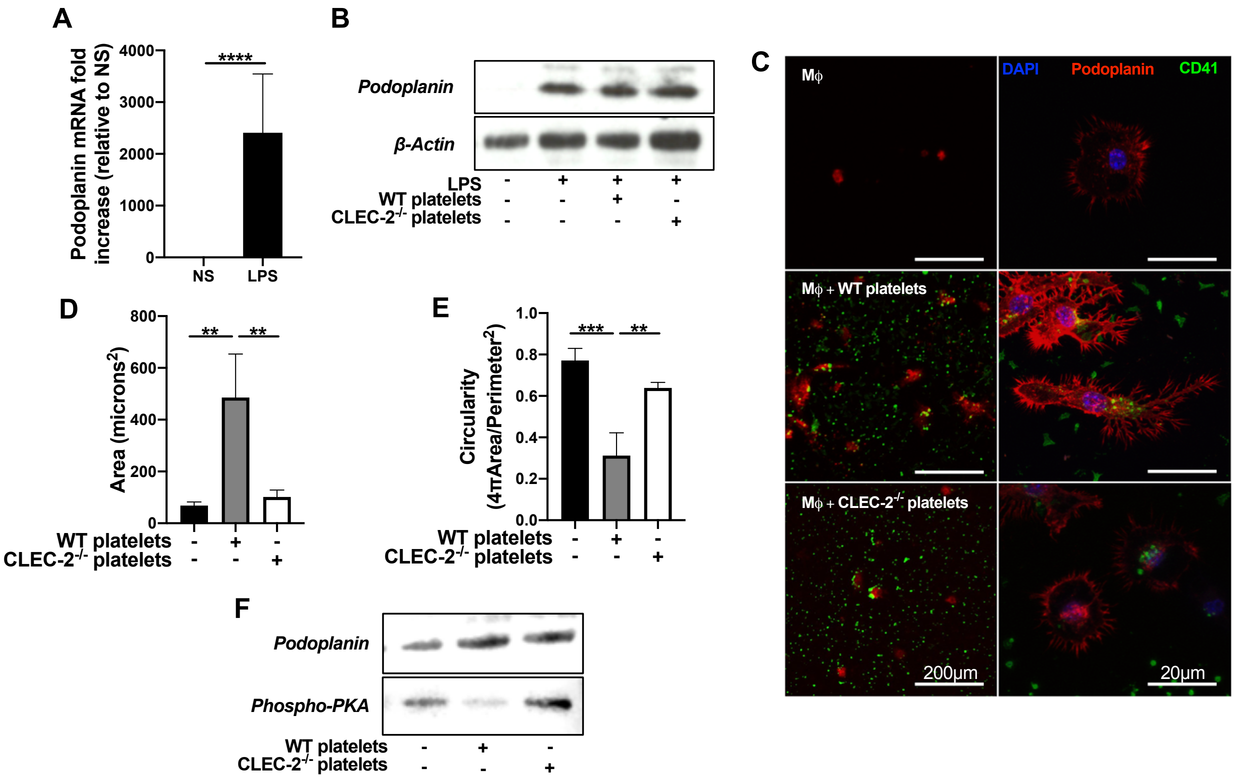


**Platelet CLEC-2 upregulates the expression of podoplanin on RAW264.7 cells and induces spreading through serines**

**(A)** RAW264.7 cells were LPS-stimulated (1μg/ml) for 24h (Mϕ). Podoplanin transcription quanitifed by qPCR from RNA. **(B)** Total podoplanin levels were assessed in Mϕ lysates incubated in the presence or absence of WT or CLEC-2-deficient platelets by western blot. Western blot are representative of 3 independent experiments. **(C)** Mϕ cells were cultured on glass and WT or CLEC-2-deficient platelets added for 1h. Cells were fixed, permeabilised and nuclei (Hoechst 33342, blue), podoplanin (red) and platelets (CD41, green) were detected using confocal microscopy. **(D)** Cell area and **(E)** circularity were analysed using ImageJ (n=3). **(F)** Podoplanin was immunoprecipitated with anti-podoplanin antibody (8.1.1), and immunostained with anti-phospho-PKA substrate antibody. Representative western blot of 3 independent experiments. The statistical significance between 2 groups was analyzed using a student's paired t-test and the statistical difference between multiple groups using one-way ANOVA with Tukey's multiple comparisons test. *$p < 0.05$ **$p < 0.01$ ***$p < 0.001$ ****$p < 0.0001$.

### Figure S3.

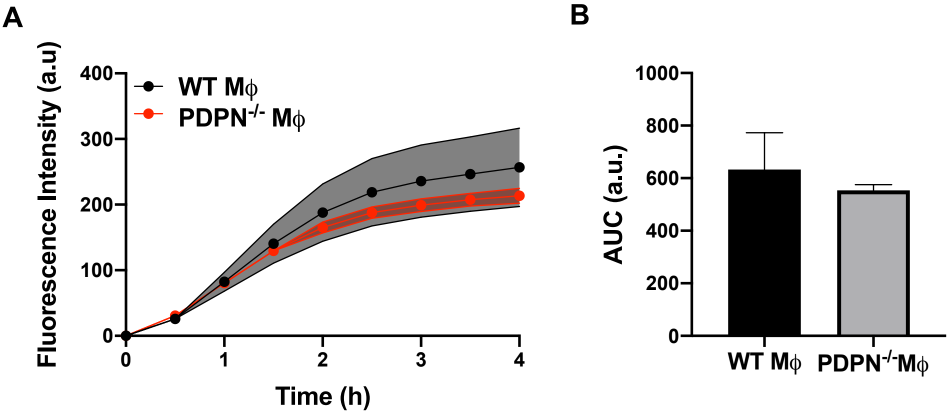


**Podoplanin deficiency does not reduce BMDM phagocytic capacity**

**(A)** WT or PDPN$^{fl/fl}$VAV Cre+ BMDMs were stimulated with LPS (1µg/ml) for 24h. **(A, B)** Phagocytosis of pH sensitive Alexa Fluor-488 conjugated *Escherichia Coli* bioparticles (3x$10^6$ beads/condition) was visualised for 4h and analysed using Incucyte SX-5 Live-Cell microscopy. (n=4). The statistical significance between 2 groups was analyzed using a student's paired t-test.

### Figure S4.

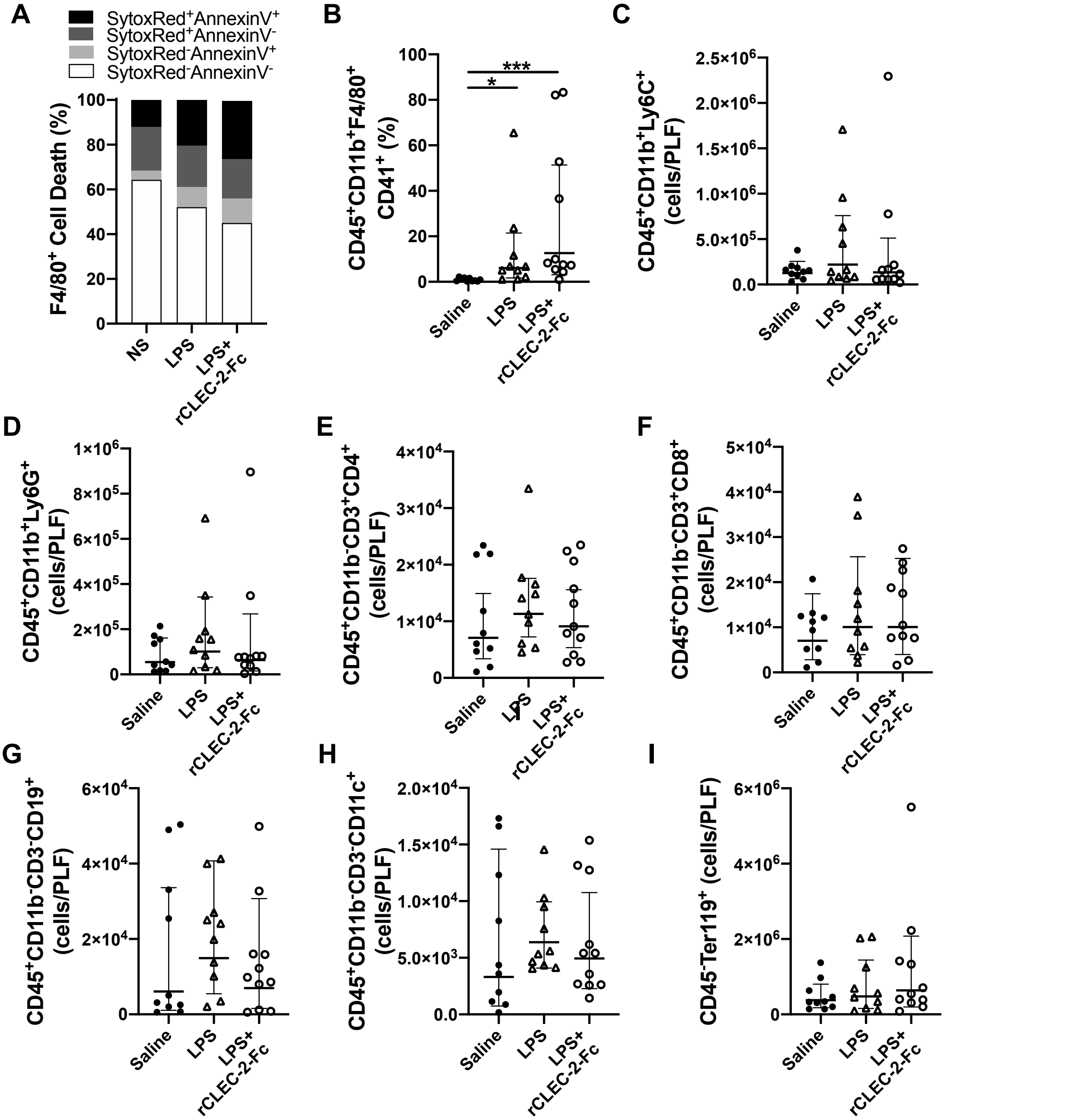


**Supplementary 4: rCLEC-2-Fc does not significantly alter macrophage death or platelet-macrophage complexes in the inflamed peritoneum**

WT mice were intraperitoneally injected with LPS (10mg/kg) for 18h followed by rCLEC-2-Fc or IgG isotype control (100µg/mouse) for and additional 4h (n=11). **(A)** Peritoneal macrophage ($CD45^+CD11b^+F4/80^+$) viability was quantified using SytoxRed and AnnexinV staining. **(B)** The percentage of platelet-macrophage complexes in the peritoneum ($CD45^+CD11b^+F4/80^+CD41^+$), total count of **(C)** monocytes ($CD45^+CD11b^+Ly6C^+$) and **(D)** neutrophils ($CD45^+CD11b^+Ly6G^+$) in the peritoneum were measured by flow cytometry. Total count of **(E)** CD4+ and **(F)** CD8+ T cells ($CD45^+CD11b^-CD3^+CD8^+$), **(G)** B cells ($CD45^+CD11b^-CD3^-CD19^+$), **(H)** dendritic cells ($CD45^+CD11b^-CD3^-CD19^-CD11c^+$) and **(I)** erythrocytes ($CD45^-Ter119^+$). The statistical significance was analyzed using a Kruskal-Wallis test. $^*p < 0.05$ $^{**}p < 0.01$ $^{***}p < 0.001$.

#### Figure S5.

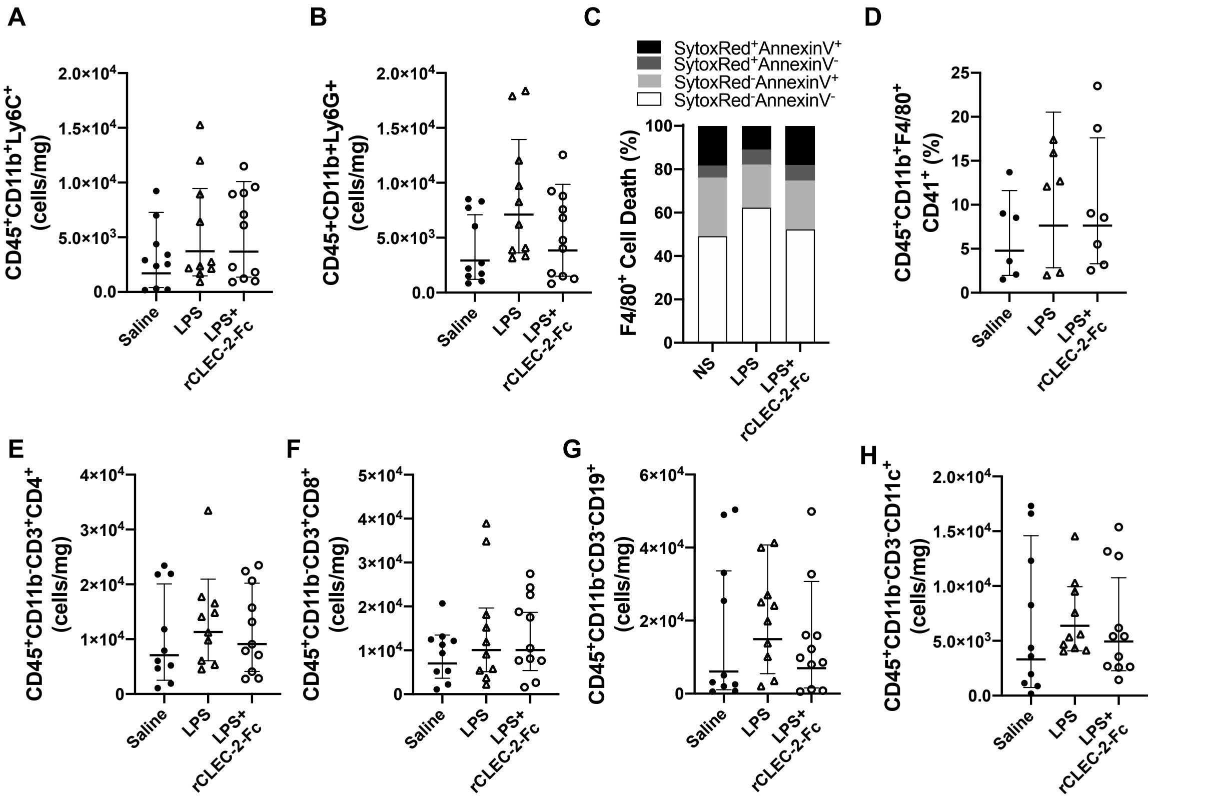


**Supplementary 5: rCLEC-2-Fc does not significantly alter splenic macrophage death or platelet-macrophage complexes**

WT mice were intraperitoneally injected with LPS (10mg/kg) for 18h followed by rCLEC-2-Fc or IgG isotype control (100µg/mouse) for an additional 4h (n=11). Total count normalised to spleen weight of **(A)** monocytes ($CD45^+CD11b^+Ly6C^+$) and **(B)** neutrophils ($CD45^+CD11b^+Ly6G^+$). **(C)** Macrophage ($CD45^+CD11b^+F4/80^+$) viability was quantified using SytoxRed $^{and}$ AnnexinV staining. **(D)** The percentage of platelet-macrophage complexes in the spleen ($CD45^+CD11b^+F4/80^+CD41^+$). Total cell counts normalised to spleen weight of **(E)** CD4+ T-Cells ($CD45^+CD11b^-CD3^+CD4^+$), **(F)** CD8+ T cells ($CD45^+CD11b^-CD3^+CD8^+$), **(G)** B cells ($CD45^+CD11b^-CD3^-CD19^+$), and **(H)** dendritic cells ($CD45^+CD11b^-CD3^-CD19^-CD11c^+$). The statistical significance was analyzed using a Kruskal-Wallis test. *$p < 0.05$ **$p < 0.01$ ***$p < 0.001$.

#### Table 1.

| Antibody (clone) | Host/Conjugate | Reference | Source |
|---|---|---|---|
| Podoplanin (8.1.1) | PE | 127408 | BioLegend |
| Podoplanin (8.1.1) | Syrian Hamster | 14-5381-85 | eBioScience |
| CD44 (IM7) | AF647 | 103018 | BioLegend |
| CD44 (IM7) | Rat | 70-0441-U100 | Tonbo Bioscience |
| CD41 (MWReg30) | PE | 12-0411-82 | invitrogen |
| CD41 (MWReg30) | BV421 | 133912 | BioLegend |
| CD41 (MWReg30) | FITC | 133904 | BioLegend |
| CD41 (MWReg30) | Rat | 553847 | BD Bioscience |
| ERM | Rabbit | 3142S | Cell Signalling |
| CD45 (30-F11) | APC | 20-0451-U100 | Tonbo Bioscience |
| CD45 (30-F11) | APC-Cy7 | 47-0451-82 | Invitrogen |
| CD11b (m1/70) | Biotin | 13-0112-82 | eBioScience |
| F4/80 (BM8) | eFluor450 | 48-4801-82 | eBioScience |
| F4/80 (BM8) | Rabbit | ab100790 | abcam |
| F4/80 (BM8) | APC | 17-4801-82 | eBioScience |
| CLEC-2 | FITC | - | internally sourced |
| Ly6C (HK1.4) | Rat | 128002 | Biolegend |
| Ly6C (HK1.4) | PERCP-Cy5.5 | 45-5932-82 | eBioScience |
| Ly6G (1A8) | V450 | 560603 | BD Horizon |
| CD80 (16-10A1) | FITC | 35-0801-U500 | Tonbo Bioscience |
| β-Actin (polyclonal) | Rabbit | ab8227 | abcam |
| Phospo-PKA (RRXS*/T* | Rabbit | 9624S | Cell Signalling |
| AnnexinV | FITC | 556420 | BD Pharmingen |
| SytoxRed | TxRed | S34859 | ThermoFisher |
| CD3 (145-2C11) | PE-Cy7 | 25-0031-82 | eBioScience |
| CD4 (RM4-5) | PE-Cy5.5 | 35-0042-82 | eBioScience |
| CD8a (53-6.7) | PE | 12-0081-82 | eBioScience |
| CD11c (N418) | FITC | 11-0114-82 | eBioScience |
| Ter119 (TER119) | Rat | 116202 | Biolegend |
| CD16/32 (93) | Rat | 14-0161-86 | eBioScience |
| CD19 (eBio1D3) | APC-Cy7 | 47-0193-82 | eBioScience |
| LYVE1 (polyclonal) | Rabbit | ab14917 | abcam |
| CCL21 (polyclonal) | Biotinylated | BAF457 | R&D |

**Antibodies used to acquire protein quanitification by flow cytometry and microscopy**
